# Maternal antiviral history synergizes with pregnancy and lactation to transfer intergenerational systemic immunity through IgG in milk

**DOI:** 10.64898/2026.08.14.744935

**Authors:** Ka Neng Cheong, Juan Sebastian Jara, Leigh M. Sewall, Stanislav Dikiy, Xuan Le, Nicole Wolman, Andrew B. Ward, R. Luke Wiseman, Alejandra Mendoza

## Abstract

Maternal immune transfer is essential for early-life health, yet whether immune experiences before pregnancy shape maternal physiology to optimize immunity in subsequent offspring is unclear. Here, we show that respiratory viral infection before pregnancy confers robust protection against lethal neonatal influenza through antibodies transferred *postpartum* in milk. Despite the predominance of IgA in milk, antiviral IgG is indispensable for protection. Pregnancy amplifies pre-existing antiviral B cell responses, while prior infection durably reprograms the mammary gland to promote transfer of circulating antiviral IgG into milk. These antibodies retain their epitope specificity, are enriched for broadly protective influenza epitopes, remain functional after passage through the neonatal intestine, and enter offspring circulation through FcRn to provide protection beyond weaning. Natural transmission of virus from infected offspring to mothers establishes maternal immunity that protects future offspring, revealing a coordinated adaptive program that links maternal immune history, pregnancy, and lactation to optimize intergenerational immunity.

GRAPHICAL ABSTRACT

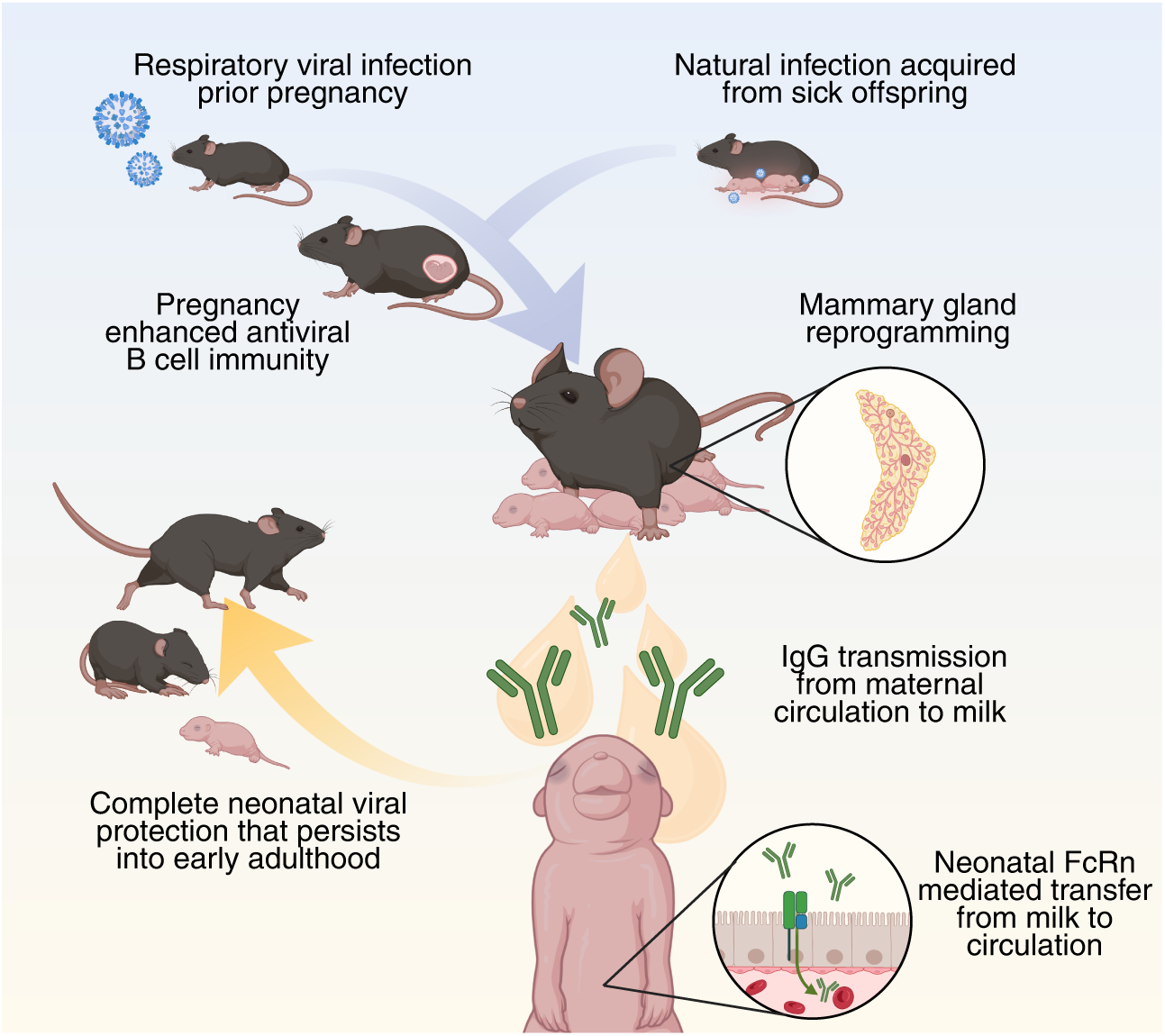

**HIGHLIGHTS:**

- Preconceptual maternal intranasal influenza infection confers complete neonatal B cell mediated protection against lethal neonatal influenza infection that persists beyond weaning into early adulthood.
- Protection can be transmitted postnatally through milk and is fully dependent on maternal IgG.
- Pregnancy enhances rather than suppresses antiviral B cell programs in the mother.
- Milk IgG targets a restricted set of conserved influenza Hemagglutinin epitopes, suggesting selective transfer of broadly protective antibody populations.
- Protective IgG in milk derives from maternal circulation, not from local B cell mammary gland production.
- Respiratory infection before pregnancy induces long-lived vascular, stromal, and epithelial transcriptional remodeling of the mammary gland.
- Neonatal Fc Receptor (FcRn) mediated transport of milk IgG into circulation is required for protection.
- Infected neonates transmit virus back to mothers, extending protection to subsequent litters for multi-generation protection.

## INTRODUCTION

Newborns are especially vulnerable to viral respiratory infections due to their developing immune systems, characterized by unique functional properties suited to the induction of tolerance, rapid innate-like responses and limited adaptive immune capacity ^1^. These immunological conditions render traditional vaccination strategies ineffective for infants under six months of age, necessitating alternative protective mechanisms ^2–4^. Accordingly, mammals have evolved to bridge this vulnerable window through the passive transfer of maternal immunity, a critical yet mechanistically poorly understood defense during early life. Passive maternal immunity refers to the vertical transfer of maternal immune factors, such as cytokines, antibodies and immune cells, to offspring during gestation and after birth via breastfeeding, providing protection against pathogens in early life and influencing neonatal immune maturation ^5,6^. From gestation through lactation, this dynamic process is shaped by maternal immune adaptations that balance fetal tolerance with antimicrobial defense ^7–10^. Consistent with this, while pregnancy is associated with increased susceptibility to certain infections, it can still support effective vaccine-induced immunity and protect newborns against pertussis and diphtheria ^11,12^. To account for this evolving immune landscape, defined windows exist during which certain immunizations are recommended or contraindicated. However, given the ethical and biological constraints of studying immunity during pregnancy, these windows are based on limited experimental data and remain incompletely understood ^13–15^.

Considering these limitations, immune exposures preceding pregnancy, or those that contribute to immunity during breastfeeding, may represent more flexible and potentially advantageous windows for shaping maternal and neonatal protection. In this context, breastfeeding serves as a key postnatal pathway for the transmission of immunity, as milk contains a diverse array of immune components including antibodies, cytokines, immune complexes, antimicrobial factors, and maternal immune cells that can influence neonatal immune development and protection ^6^. Beyond playing a role in passive protection through milk, immune populations, including macrophages, eosinophils and intraepithelial lymphocytes have been shown to drive mammary epithelial differentiation, tissue remodeling, and milk production, demonstrating that the mammary gland is an immune-regulated organ ^16–20^. Together, these observations highlight a bidirectional relationship in which maternal immunity can both transfer protection to offspring and program the functional capacity of the mammary gland itself. Therefore, maternal immune history, particularly exposures sustained prior to pregnancy, is a critical and underexplored determinant of mammary gland biology and neonate protection against infection in early life.

While it is well established that infections induce durable changes in immune composition and function across tissues ^21–24^, whether such exposures influence mammary gland development, lactational output, or the quality and magnitude of immunity transferred to neonates is currently unknown. In this regard, respiratory viral infections elicit robust systemic and mucosal immune responses that could reshape immune populations recruited to the mammary gland during pregnancy and lactation. However, studies examining infection in this setting have largely focused on pathogen transmission through milk or on infections localized to the gland itself, such as mastitis, rather than on how prior infections at distal sites imprint long-term changes on mammary gland function and induction of neonatal protection ^25^. Here, we asked if maternal respiratory viral infection prior to pregnancy affects the mammary gland to promote the transmission of protective immunity in offspring. We demonstrate that maternal immune mucosal priming enhances the vascular and structural remodeling gene programs in the mammary gland, allowing for the transfer of protective immunity to offspring against lethal influenza neonatal respiratory infection. We show that this maternal protection can be transferred through nursing and is fully dependent on maternal IgG antibodies. These findings establish insights into maternal-infant immune protection, which can inform potential prophylaxis strategies for advancing maternal and pediatric health.

## RESULTS

### Preconceptual maternal respiratory infection protects offspring against lethal influenza infection

Neonates are highly susceptible to respiratory viral infection. To model this neonatal susceptibility, we infected neonates at postnatal day 5 (P5) intranasally with PR8 influenza A virus (IAV). Neonatal IAV infection (2 Focus-forming units (FFU)) caused severe disease by 7-10 days, resulting in 95.7% mortality before weaning age (Fig. 1A). Using this lethal neonatal infection model, we examined whether maternal immunity primed by natural infection before pregnancy could protect offspring during this vulnerable period. We intranasally infected female mice 18-21 days prior to mating with a sublethal dose of IAV (1FFU). In adults, this dose, which is approximately one-tenth of the Median Lethal Dose (LD_50_), caused infection without overt disease as it induced robust T cell and B cell responses with insignificant weight loss and no change in fecundity (Fig. 1B-G, S1A-B, S2, S3A-E). We found that neonates born to IAV primed females (42-47 days after maternal priming) uniformly survived lethal IAV infection at P5 and reached weaning age with minimal to undetectable viral transcripts in their lungs (Fig. 1F, H-I). In contrast, neonates born to females that were primed with PBS all succumbed to infection and showed IAV transcripts in the lung. This indicates that infection before pregnancy promotes robust and protective vertical immunity to neonates against respiratory viral infection.

**Figure 1.**
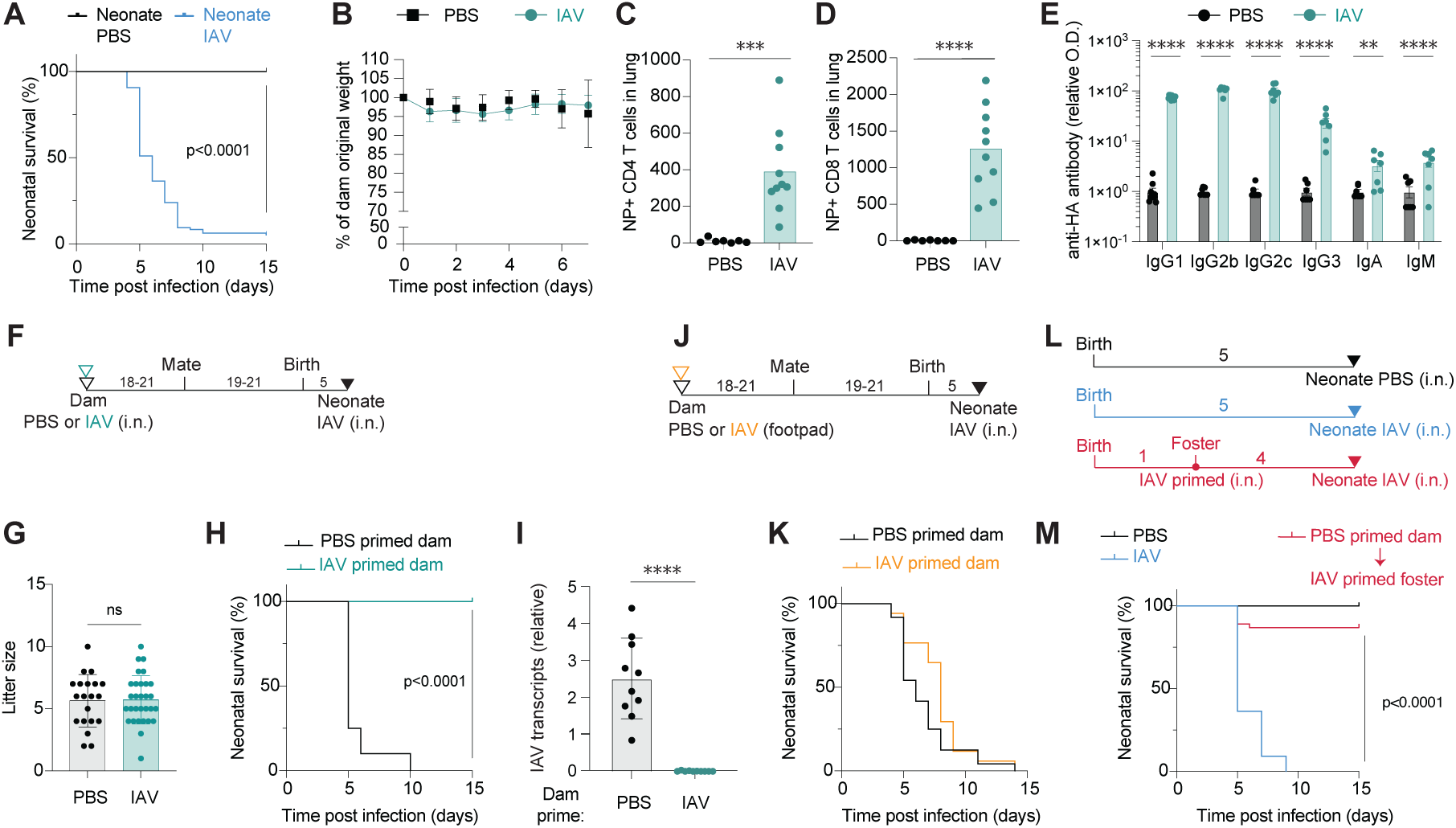
Preconceptual maternal respiratory infection protects neonates from lethal influenza infection. **(A)** Survival of neonates following intranasal infection with IAV (2 FFU, i.n.) or PBS at postnatal day 5 (P5). PBS n=26; IAV n=96. **(B-E)** Dams were infected with IAV (1 FFU, i.n.) and mated 18-21 days post infection. **(B)** Percent body weight loss after IAV infection. IAV (n=4), PBS (n=4). (**C-D**) Number of influenza nucleoprotein specific NP+ CD4+ T cells **(C)** and NP+ CD8+ T cells **(D)** in the lung of dams analyzed by flow cytometry at P5. **(E)** Relative titers of influenza hemagglutinin (HA) specific antibodies of indicated isotypes in dam serum quantified by ELISA on P5. Optical density at 450 nm (O.D.) was normalized to the average O.D. of control group per isotype. Non-detectable values were assigned a zero. **(F)** Litter size of IAV and PBS primed dams. **(G-I)** Dams were infected with IAV (1 FFU, i.n.) or PBS and mated 18-21 days post infection. Offspring of IAV or PBS primed dams were infected (2 FFU, IAV i.n.) at P5. **(G)** Schematic of experimental setup. **(H)** Percent survival of neonates following IAV infection from indicated dams. PBS n=20; IAV n=74. **(I)** IAV viral RNA transcripts in lungs of IAV challenged neonates born to PBS or IAV primed dams, quantified 3 days post neonatal infection by RT-qPCR relative to lung *Actb* transcripts. Non-detectable values were assigned a zero. **(J-K)** Dams were injected with IAV (50 FFU) or PBS in the footpad and mated 18-21 days post inoculation. Offspring of IAV or PBS inoculated dams were infected with IAV (2 FFU, i.n.) at P5. **(J)** Schematic of experimental setup. (**k**) Survival of neonates from indicated dams following neonatal IAV infection at P5. PBS n=24; IAV n=17. **(L-M)** Foster dams were infected with IAV (1 FFU, i.n.) and mated 18-21 days post infection. For cross fostering, neonates from naïve dams were transferred to foster cage by P1. Neonates from naïve dams and cross-fostered were infected with IAV (2 FFU, i.n.) or PBS on P5. **(L)** Schematic of experimental groups. **(M)** Survival of neonates following IAV infection at P5. PBS n=8; IAV n=11; PBS-primed dam with IAV-primed foster dam n=46. Dots represent data from individual mice, bars show means, error bars show SD (**C, D, E, G** and **I**). Statistical significance determined by two-tailed unpaired t-test with Welch’s correction (**C, D, G,** and **I**), by 2-way ANOVA with Sidak’s multiple comparisons test (**E**) or using survival Logrank test (**A, H, K** and **M**). *p<0.05, ** p<0.01, *** p<0.001, **** p<0.0001.

Mucosal and systemic routes of infection engage distinct immune compartments and effector programs, resulting in differences in the quality of local versus systemic protection ^26,27^. To test if distinct routes of immune priming influence vertically transferred protection to offspring, we performed footpad priming of female mice with a high dose of IAV (50 FFU, 50-fold higher than the dose used for intranasal priming). Primed females were mated 21 days after footpad immunization and their pups were then challenged with the neonatal IAV lethal dose (2 FFU) at P5. Despite footpad inoculation inducing influenza specific T cell and B cell responses, all neonates born to footpad primed females succumbed to lethal IAV challenge at similar rates to those born to mock primed (PBS) females (Fig. 1J-K, S3F-J). Of note, intranasal IAV exposure, even at a 50-fold lower dose, generated a stronger influenza specific response compared to footpad priming (Fig. S3K-L). These data suggest that respiratory mucosal viral exposure is superior to peripheral priming in inducing maternal transfer of protective immunity to neonates, potentially owing to the route of exposure and the greater magnitude of the immune response it elicits.

The vertical transfer of maternal immune factors can proceed postnatally through nursing, raising the possibility that lactational transfer alone may be sufficient to induce the neonatal protection conferred by maternal IAV exposure. To test this, we performed fostering experiments in which neonates born to unprimed females were fostered by IAV primed females that were infected 47 days before neonatal transfer (1 FFU). Within 24 hours of birth, pups were transferred and 4 days post transfer (at P5) they were challenged with a lethal neonatal IAV dose (Fig. 1L-M). Neonates born to naïve females and nursed by IAV primed fosters showed 87% survival by weaning age, indicating that nursing alone is sufficient to provide near complete protection against lethal IAV infection (Fig. 1M). Together, these data demonstrate that preconceptual respiratory infection induces a durable and route-dependent maternal immune response that is sufficient to protect offspring against IAV, and that this protection is efficiently transmitted postnatally through nursing.

### Maternal IgG antibodies transferred through milk can confer protection against neonatal viral infection

Having established that protective immunity against respiratory IAV infection can be transferred postnatally via breastfeeding, we next asked whether antibodies are required for this protection. Among the factors transferred through milk, antibodies produced by maternal B cells are compelling candidates, as they can provide immediate defense against pathogens while also shaping neonatal immune responses during their critical early window of development ^6^. Maternal immunoglobulins in milk have been shown to be protective in the context of enteric diseases and gut microbiome regulation ^28–31^, but the potential of maternal derived milk antibodies for protecting neonates to respiratory viral infection is poorly understood. To determine whether maternal B cell responses are necessary for neonatal protection, we infected female *μMT^⁻/⁻^* mice, which lack mature B cells, intranasally with IAV (1 FFU) 21 days before mating. These mice were bred with wild-type males to generate phenotypically wild-type offspring only lacking maternal B cell-derived immunity. Neonates from these females were challenged with a lethal dose of IAV at P5. All neonates born to *μMT^-/-^*female mice were susceptible to lethal IAV infection regardless of maternal priming status (Fig. 2A-B). We next asked whether B cells were required for the protective effects mediated specifically by nursing. To address this, we fostered neonates from *μMT^⁻/⁻^* dams to IAV primed wild-type dams, generating offspring that received milk from IAV-primed B cell-sufficient mice. We found that fostering rescued neonatal mortality in IAV-infected pups born to B cell deficient dams, showing 94.7% survival (Fig. 2C-D). These data indicate that B cells are required for maternal transfer of neonatal protection against lethal IAV infection, and that milk from B cell-sufficient primed females is sufficient to confer this protection.

**Figure 2.**
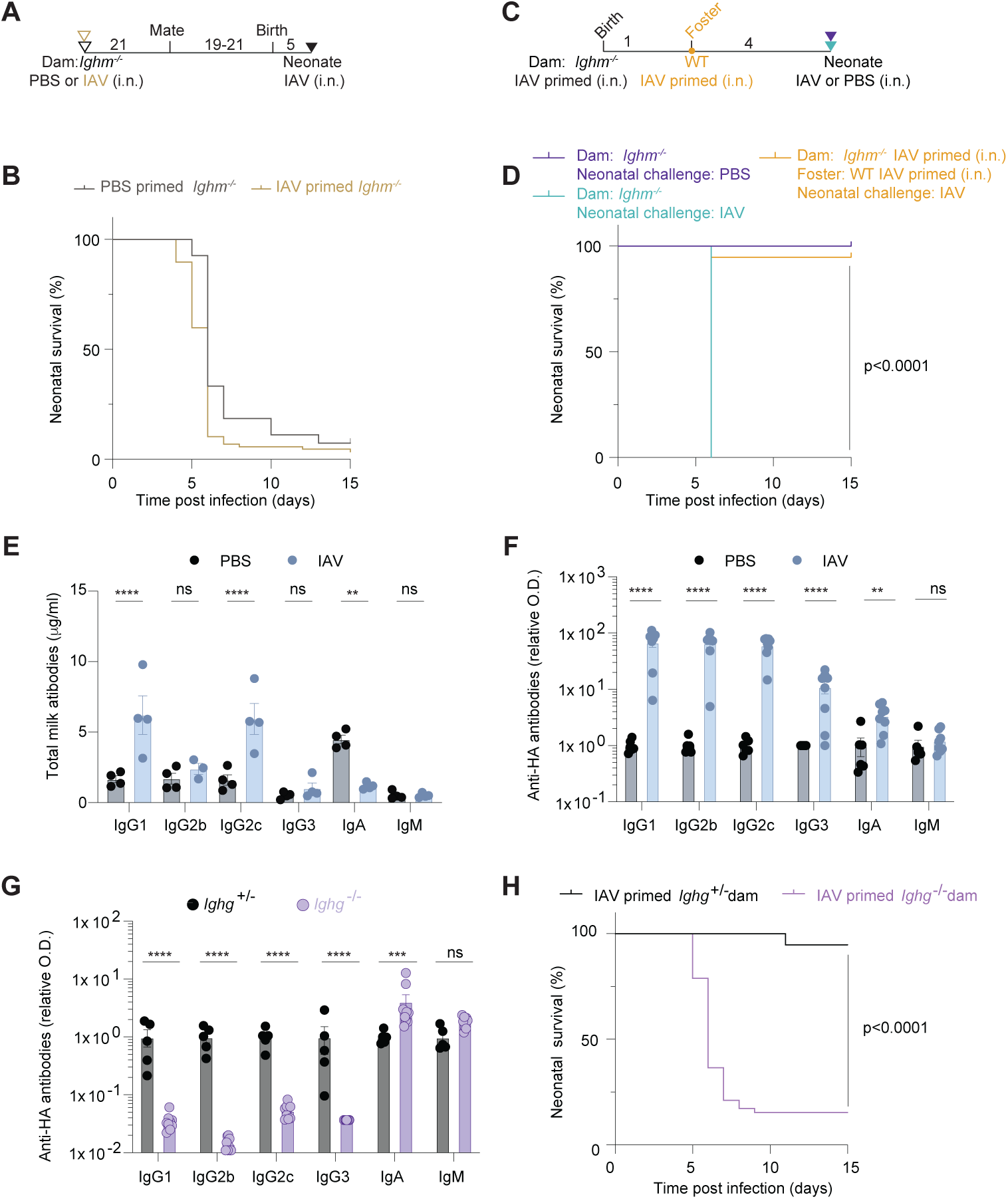
Maternal IgG antibodies transferred through milk confer neonatal protection against influenza. **(A-B)** B cell deficient *Ighm^-/-^* dams were infected with IAV (1 FFU, i.n.) or PBS and mated 18-21 days post infection. Offspring of IAV or PBS primed *Ighm^-/-^* dams were infected (2 FFU, IAV i.n.) at P5. **(A)** Schematic of experimental setup. **(B)** Percent survival of neonates following IAV infection from indicated dams. PBS primed *Ighm^-/-^*n=27; IAV primed *Ighm^-/-^* n=87. **(C-D)** *Ighm^-/-^ dams and* WT foster dams were infected with IAV (1 FFU, i.n.) and mated 18-21 days post infection. For cross fostering, neonates from IAV-primed *Ighm^-/-^* dams were transferred to foster cage by P1. Neonates from IAV-primed *Ighm^-/-^* dams and cross-fostered by WT IAV-primed dams were infected with IAV (2 FFU, i.n.) or PBS on P5. **(C)** Schematic of experimental groups. **(D)** Survival of neonates following IAV infection at P5. PBS n=9; IAV n=7; *Ighm^-/-^* IAV-primed dam with WT IAV-primed foster dam n=38. **(E-F)** Dams were infected with IAV (1 FFU, i.n.) and mated 21 days post infection and antibodies were quantified in milk at P5 by ELISA. **(E)** Total antibody concentration of indicated isotypes. **(F)** Relative titers of HA-specific antibodies of indicated isotypes. Optical density at 450 nm (O.D.) was normalized to the average O.D. of control group per isotype. Non-detectable values were assigned a zero. **(G-H)** *Ighg^-/-^* and littermate control (*Ighg^+/-^)* dams were infected with IAV (1 FFU, i.n.) and mated 21 days post infection. **(G)** Relative titers of HA specific antibodies of indicated isotypes 14 days post infection in serum quantified by ELISA. Optical density at 450 nm (O.D.) was normalized to the average O.D. of control group per isotype. Non-detectable values were assigned a zero. **(H)** Offspring of *Ighg^-/-^*and littermate control (*Ighg^+/-^)* dams were infected with IAV (2FFU, i.n.) on P5. Percent survival of neonates following IAV infection. IAV primed *Ighg^+/-^*dam n=38; IAV primed *Ighg^-/-^* dam n=52. Dots represent data from individual mice, bars show means, error bars show SEM (**E-G)**. Statistical significance determined by 2-way ANOVA with Sidak’s multiple comparisons test **(E-G)**, or using survival Logrank test **(B, D, H)**. ns= not significant; *p<0.05, ** p<0.01, *** p<0.001, **** p<0.0001.

The dependence on B cells for neonatal protection, combined with the sufficiency of milk to transfer this protection, suggested that maternally derived influenza-specific antibodies may mediate neonatal defense against lethal IAV infection (Fig. 2A-D). Although IgA antibodies constitute the majority of the antibodies present, IgM and IgG antibodies are also found in milk ^32^. Moreover, maternal immune responses have the potential of altering the isotypes and specificity of antibodies found in milk. Indeed, analysis of human breast milk from individuals previously infected with or vaccinated against respiratory viruses have been shown to contain virus-specific IgA and IgG antibodies ^33–36^. Therefore, we profiled the antibody isotypes present in milk collected at P5 from IAV- and mock-primed female mice. We found that milk from IAV primed female mice had a shift in overall composition of antibody isotypes with more IgG1 and IgG2c antibodies than PBS-primed females and an overall decrease in IgA (Fig. 2E). The increase in IgG1 and IgG2c was also evident in maternal serum, although we observed no change in total IgA (Fig. S4A). Antigen specific antibodies to influenza hemagglutinin (HA) were present in IAV primed female serum, confirming that 47 days post infection females still had detectable antigen specific antibodies (Fig. 1E). HA-specific antibodies were also detected in milk collected from IAV-primed females at P5, and were below the level of detection in milk from PBS-primed females (Fig. 2F). Notably, the HA-reactive antibody response in milk predominantly comprised IgG, with minimal HA-specific IgA detected (Fig. 2F). Consistent with this, HA-specific IgG persisted in milk sampled at P11, whereas HA-specific IgA levels further declined, indicating that the IgG dominance was not simply a reflection of IgG-rich composition in early milk (Fig. S4B). Together, these findings demonstrate that maternal IAV exposure results in the transfer of antigen-specific antibodies to milk, with the response heavily polarized toward IgG.

Given the abundance of IAV-specific IgG antibodies present in milk, we sought to directly test whether maternal IgG antibodies are required for neonatal protection. For this, we evaluated neonates born to pre-conceptually primed IgG-deficient female mice (*Ighg^-/-^*), which lack all IgG isotypes (IgG1, IgG2b, IgG2c, and IgG3)^28^. *Ighg*^-/-^ and littermate control (*Ighg^+/-^*) female mice were infected with IAV (1FFU). As influenza viral clearance during primary infection is not dependent on B cells^37^, we found that *Ighg^-/-^* mice showed viral clearance similar to that observed in WT mice, with undetectable viral load observed at day 14 post infection (Fig. S4C). As expected*, Ighg^-/-^* females produced no HA-specific IgG antibodies but were able to generate influenza-specific IgA and IgM, both of which were present at elevated levels compared to IgG-sufficient controls (Fig. 2G, S4D). We bred these mice 21 days post infection to wild type males to ensure progeny were phenotypically wild-type. Neonates born to IAV primed *Ighg^-/-^*and littermate control (*Ighg^+/-^*) female mice were infected at P5 with a lethal dose of IAV (2FFU). All neonates from *Ighg^-/-^* dams succumbed to lethal IAV infection, while those born from *Ighg^+/-^*littermate controls were fully protected (Fig. 2H). Collectively, these data demonstrate that neonatal protection against lethal respiratory IAV infection is mediated by IgG secreted from maternal B cells that is transferred to neonates through breastfeeding. Moreover, despite compensatory elevation of IgA and IgM in IgG-deficient dams, these isotypes are insufficient to substitute for IgG, establishing maternal IgG as the critical mediator of milk-derived neonatal immunity against influenza.

### Pregnancy enhances antiviral B cell responses established before conception

The requirement for IgG in mediating neonatal protection, suggests that pregnancy must preserve or even enhance antiviral IgG responses to enable effective transfer to the offspring. However, as maternal IgG antibodies can cause severe pathology when directed against fetal antigens (e.g., anti-RhD IgG antibodies in hemolytic disease and fetal/neonatal alloimmune thrombocytopenia^38^), pregnancy must carefully balance the need to maintain protective IgG responses with the danger of IgG-mediated fetal harm. This raises the important question of how pregnancy affects a pre-existing B cell response induced by prior viral infection. To determine whether pregnancy alters established antiviral immune responses, we compared influenza-specific immunity in nulliparous and parous females. Female mice were infected intranasally with IAV and, 21 days later, were either mated or left unmated. Antiviral T and B cell responses were analyzed 45-47 days after infection, when females that had been mated were actively lactating. In the lung-draining mediastinal lymph nodes (medLN), influenza-specific CD4⁺ and CD8⁺ T cell responses were induced to a similar extent in nulliparous and lactating females relative to their respective uninfected controls (Fig. S2, S5A-B). However, lactating females exhibited a modest but significant increase in T_H_1 cells marked by expression of the transcription factor T-bet compared to nulliparous mice (Fig. S5C-D). By comparison, antiviral B cell responses were markedly enhanced following pregnancy. Lactating dams exhibited increased frequencies and absolute numbers of germinal center (GC) B cells in the medLN compared with nulliparous females (Fig. 3A-B). Moreover, GC B cells from lactating dams showed a selective increase in T-bet expression (Fig. 3C), a key regulator of antiviral B cell immunity shown to be a hallmark of antiviral responses required for switching to IgG2, the dominant antiviral IgG isotype in mice ^39–47^. The selective increase in T-bet+ GC B cells during lactation suggests that the gestational environment amplifies the B cell program required for protective IgG2c production, consistent with the enhanced antibody transfer and neonatal protection observed in our model. Indeed, despite the physiological reduction in total serum IgG concentration that accompanies pregnancy and lactation as a consequence of plasma volume expansion^48^, HA-specific IgG2c antibodies were significantly enriched within the circulating antibody pool of lactating dams compared to nulliparous females (Fig. 3D, S5E). Thus, pregnancy selectively enriches protective antiviral antibodies despite the hemodilution that accompanies gestation and lactation.

**Figure 3.**
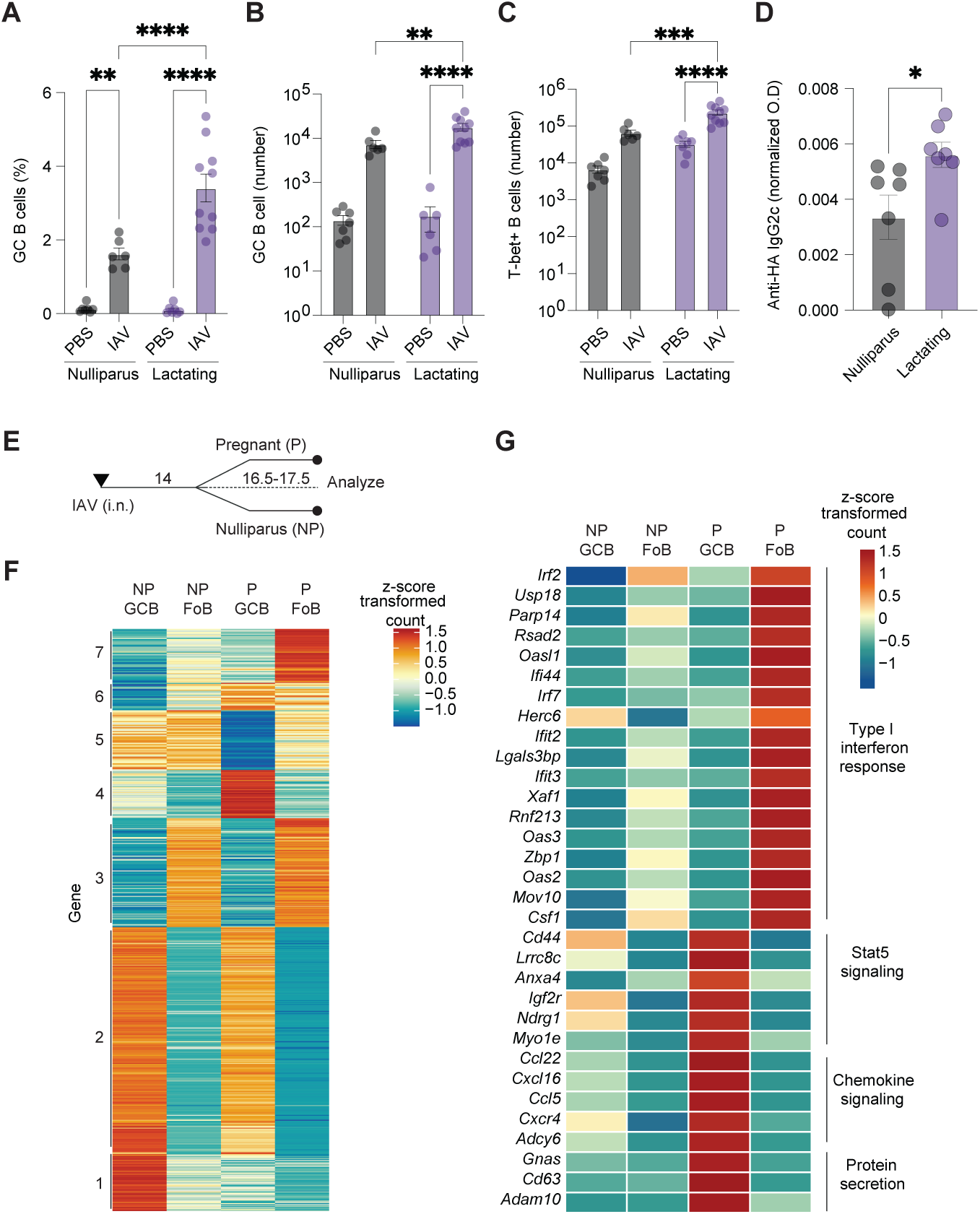
Antiviral B cell programs established before conception are enhanced during pregnancy and lactation. **(A-C)** *Tbx21^TdTomato-Cre^* mice were infected with IAV (1 FFU, i.n.) or PBS and mated or left unmated 21 days post infection. B cells were analyzed by flow cytometry at P5 from medLN following CD45 I.V. labeling. **(A)** Frequency of GC B cells of total B cells. **(B)** Cell numbers of GC B cells and **(C)** Tbet+ total B cells. **(D)** Serum from nulliparous and lactating dams were analyzed for HA-specific IgG2c antibodies by ELISA on P5. Optical density at 450 nm (O.D.) was normalized to the average IgG2c concentration of the same samples of each group. **(E-G)** Dams were infected with IAV (1FFU, i.n.). Following 14 days post infection dams were mated or left unmated. Germinal center (GC) and follicular (FoB) B cells were sorted from the mediastinal LNs of nulliparous (NP) and pregnant (P) mice at E16.5-E17.5 and analyzed by RNA-seq. **(E)** Schematic of experimental setup. **(F)** Heatmap showing k-means clustering of differentially expressed genes across all pairwise comparisons. Values are z-score normalized by row. **(G)** Expression patterns of select genes associated with interferon response, Stat5 signaling, and protein secretion. Dots represent data from individual mice, bars show means, error bars show SEM **(A-D)**. Statistical significance was determined by 2-way ANOVA with Sidak’s multiple comparisons test **(A-C)** or two-tailed unpaired t-test with Welch’s correction **(D)**. *p<0.05, ** p<0.01, *** p<0.001, **** p<0.0001. **(E-G)** RNA-seq analysis was performed on three biological replicates.

To gain further insight into how pregnancy reshapes antiviral B cell function, we performed transcriptional profiling of naïve and GC B cell subsets sorted from the lung-draining mediastinal lymph nodes of IAV-primed nulliparous or pregnant (E16.5-17.5) females 30 to 31 days following infection (Fig. 3E-G). The corresponding populations were defined, sorted, sequenced, and analyzed as nulliparous naïve, pregnant naïve, nulliparous GC, and pregnant GC B cells. By examining shared and distinct gene expression patterns across these four populations, we identified groups of genes that were uniquely, differentially, or commonly expressed, revealing that overall pregnancy enhanced several aspects of antiviral immunity (Fig. 3F). Specifically, in naïve B cells, pregnancy induced strong upregulation of interferon (IFN) α and γ response genes, including *Irf2, Irf7, Oas2, Oas3, Oasl1, Ifit2, Ifit3* and *Csf1* (Fig. 3G). This interferon primed transcriptional state suggests that naïve B cells during pregnancy are poised for enhanced responses to viral antigens. In GC B cells, pregnancy exhibited minimal interferon gene induction, but displayed a coordinated program characterized by enhanced expression of genes involved in cellular migration (*Cxcl16, Ccl5, Ccl22, Cxcr4*), Stat5 regulated genes (*Cd44, Anxa4, Lrrc8c, Ndrg1*) and protein secretion signatures (*Gnas, Cd63, Adam10*). Interestingly, among the most prominent transcriptional changes was the enrichment of STAT5 target genes in GC B cells from pregnant compared to non-pregnant IAV-primed females (Fig. 3G). STAT5 is primarily phosphorylated and activated by IL-2 family interleukins (IL-2, IL-7, IL-9, IL-15), hematopoietic growth factors (erythropoietin and thrombopoietin), and prolactin, which rises dramatically during late pregnancy and lactation ^49,50^. This raises the possibility that prolactin, together with other cytokines and growth factors that activate the JAK-STAT5 axis during pregnancy, can synergize with GC B cell responses, thereby linking the hormonal state of pregnancy to the potentiation of protective antibody production required for neonatal defense. Thus, pregnancy seems to simultaneously enhance an interferon-primed state on naïve B cells and enhance effector programs on GC B cells, optimizing maternal antiviral antibody-mediated protection of the offspring. Together, these data indicate that pregnancy, during mammary gland development, enhances antiviral B cell functional programs established before conception, consistent with the increased IgG production and transfer necessary for neonatal protection.

### IgG transferred from maternal circulation to milk provides neonatal anti-viral protection

Having established that pregnancy enhances antiviral B cell transcriptional programs that support protective IgG production, we next asked whether the primed mammary gland actively facilitates antibody transfer. To this end we tested whether maternal priming prior to pregnancy alters the immune composition and functional state of the lactating mammary gland. Female mice were infected with IAV 21 days prior to mating, and the immune composition of the mammary glands were analyzed at P5. We found that total immune cells (CD45+), myeloid cells, T and B lymphocyte populations were present in the mammary gland of IAV primed dams at similar numbers to those observed in mock primed (PBS) dams (Fig. S6, S7A-G). While overall T cell numbers were comparable, we observed a selective accumulation of influenza-specific CD4⁺ and CD8⁺ T cells in the mammary glands of IAV-primed dams compared to PBS controls (Fig. S7H-I). This increase in antigen specific T cells is unlikely to be driven by viral infection in the mammary gland itself, as we did not detect viral transcripts in this tissue following infection (Fig. S7J, 6B). We also quantified plasma cells (isotypes IgG1, IgG2b, IgG2c, IgM, IgA) in the mammary glands, lungs, spleen and bone marrow of IAV or PBS primed lactating dams. While we did not detect differences of IgG producing plasma cells in mammary glands and spleens of IAV primed mice, those that were IAV primed showed increases of IgG2b+ and IgG2c+ plasma cells in lungs, as well as significantly increased IgG2c+ plasma cells in bone marrow (Fig. 4A-B, S8A-B). Thus, while prior IAV exposure induces modest changes in the overall immune composition of the mammary gland, including selective accumulation of antigen-specific T cells and a reduction in IgA+ plasma cells, we find no evidence of local accumulation of IgG-producing plasma cells within the gland itself.

**Figure 4.**
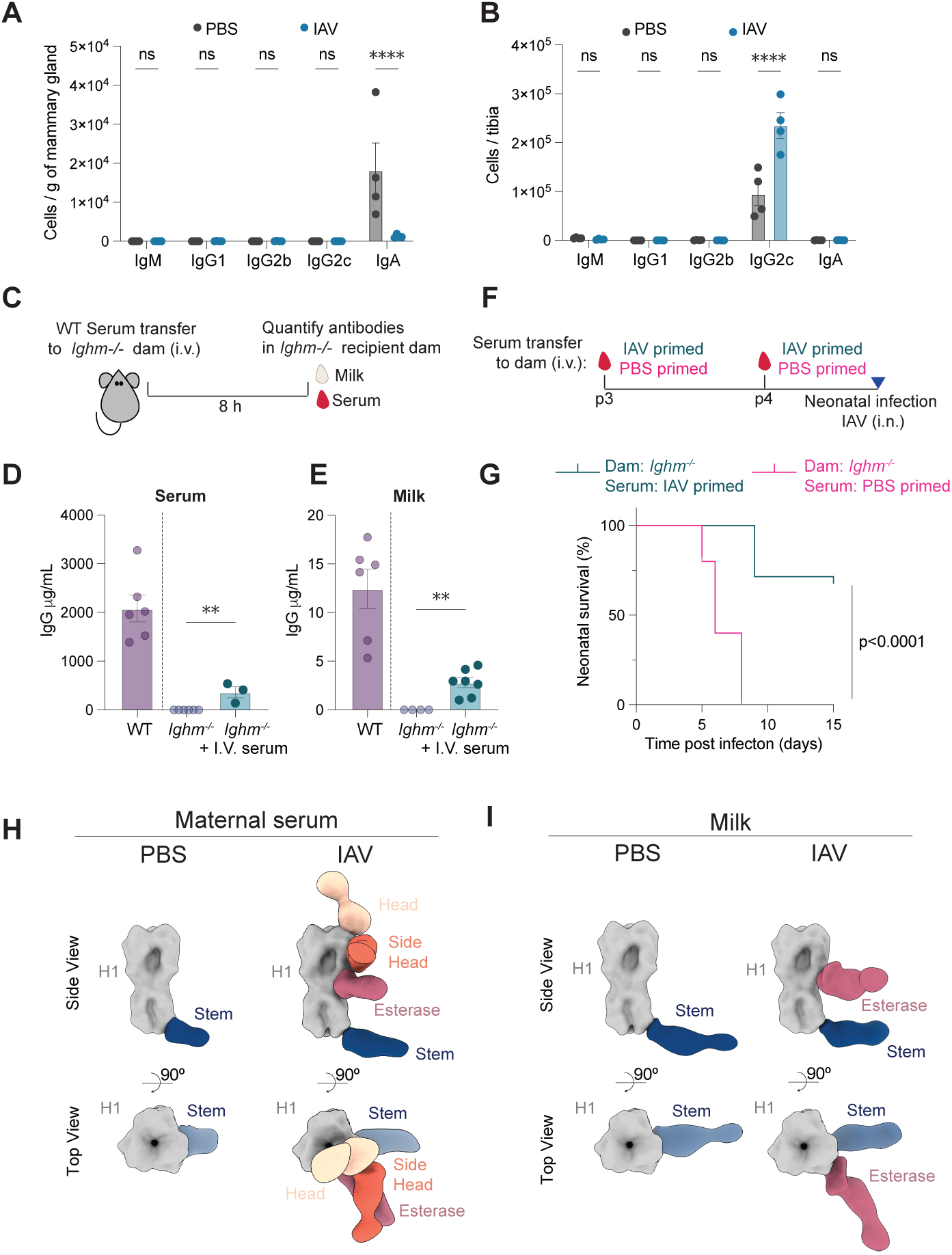
Selective transfer of IgG from maternal circulation to milk protects neonates against lethal influenza. **(A-B)** Dams were infected with IAV (1 FFU, i.n.) or PBS and mated 21 days post infection. Plasma cells were analyzed by flow cytometry at P5 from the mammary gland **(A)** and bone marrow **(B)** following CD45 I.V. labeling. **(C-E)** Serum from WT mice was collected and transferred to *Ighm^⁻/⁻^* dams at P1 or P2, and samples were collected eight hours after transfer. **(C)** Schematic of experimental setup. IgG antibody was quantified in serum **(D)** and milk **(E)** of *Ighm^⁻/⁻^* recipient dams by ELISA. **(F-G)** WT mice were infected with IAV (10 FFU, i.n.) or PBS. IAV-primed or PBS-primed serum from IAV- or PBS-primed mice was transferred to *Ighm^-/-^* dams at P3 and P4. Neonates of *Ighm^-/-^* dams receiving serum were challenged with IAV (2FFU, i.n.) at P5. **(F)** Schematic of experimental setup. **(G)** Survival of neonates nursed by *Ighm^-/-^* dams receiving IAV or PBS-primed serum. PBS primed serum n=10; IAV primed serum n=28. **(H-I)** Composite figure from EMPEM analysis by ns-EM of polyclonal antibody responses from maternal sera **(H)** and milk **(I)** collected at P5 to influenza H1 PR8 trimer, and the corresponding epitopes were mapped. Top and side views of segmented composite maps are shown, with previously identified HA epitopes head (tan), side head (orange), esterase (pink) and stem (blue) colored accordingly. The H1 PR8 trimer is shown in grey. Dots represent data from individual mice, bars show means, error bars show SEM (**A-B, D-E**). Statistical significance was determined by 2-way ANOVA with Sidak’s multiple comparisons test (**A-B, D-E**), or survival Logrank test (**G**). For EMPEM, sera and from PBS- and IAV-primed dams were pooled from 6 mice per group, milk from PBS- and IAV-primed dams were pooled from 5 mice per group **(H-I)**. ns= not significant; *p<0.05, ** p<0.01, *** p<0.001, **** p<0.0001.

The predominant accumulation of IgG producing plasma cells in the bone marrow, rather than in the mammary gland itself (Fig. 4A-B), suggested that the majority of protective IgG present in milk likely derives from maternal circulation rather than local production. To directly test this, we transferred immune serum from IAV infected (nulliparous) mice into naïve *μMT^-/-^* lactating dams. As *μMT^-/-^* females lack mature B cells and cannot produce any antibodies of their own, any immunoglobulins detected in milk from these recipients are exclusively derived from the transferred serum. IgG antibodies transferred intravenously into lactating dams were detectable in serum and milk of recipients within 8 hours (Fig. 4C-E), demonstrating efficient and rapid antibody transfer from circulation into milk. In contrast, transferred IgA was detectable in the serum of recipients but poorly detected in the milk, likely due to low concentration in the donor serum, inefficient transfer from circulation into milk, or both (Fig. S8C-D). This is consistent with IgA being predominantly produced and secreted into milk locally within the mammary gland. To determine whether this circulation-derived IgG is sufficient to confer neonatal protection, we challenged pups nursing from serum-recipient *μMT^-/-^* dams with a lethal dose of IAV at P5. Upon neonatal lethal IAV challenge, >60% pups nursed by dams receiving immune serum transfer survived infection, whereas all control pups nursed by dams receiving naïve serum succumbed to infection (Fig. 4F-G). These results suggest that circulating antibodies alone are sufficient to confer milk-mediated neonatal protection.

To characterize the polyclonal maternal antibody response elicited by IAV infection, we used electron microscopy-based polyclonal epitope mapping (EMPEM) to map the epitopes on HA trimer targeted by antibodies in both serum and milk^51–53^. Purified IgG from serum and milk of IAV-infected dams was digested into Fabs and complexed with the IAV HA trimer, and the resulting immune complexes were visualized by negative-stain EM. In the serum, polyclonal antibodies were found to target multiple sites on HA, including the head, side head, esterase, and stem regions (Fig. 4H, S9A-B). The antibody response in milk was more restricted, with antibodies predominantly recognizing the esterase and stem epitopes (Fig. 4I, S9C-D), and esterase-directed antibodies appearing substantially more abundant than in serum. Importantly, the overlap in antibody specificities between serum and milk provides independent structural support for the transfer of circulating maternal IgG into milk. Notably, the HA stem is a conserved target of broadly neutralizing influenza antibodies, suggesting that the selective transfer of these antibodies may maximize the breadth of antiviral protection conveyed to offspring. Reconstructions of the IAV trimer bound to side-head and esterase polyclonal antibodies revealed a slightly more open trimer conformation, likely reflecting antibody-induced conformational changes at these sites (Fig. 4H-I).

### Preconceptual viral respiratory infection induces long term reprograming of the mammary gland

The ready transfer of protective IgG from circulation to milk, as well as immune cell composition changes, suggest that the mammary gland itself may be remodeled to enhance antibody transfer and confer protection following IAV priming. To gain a more comprehensive and unbiased understanding of how prior IAV exposure reshapes the mammary gland, we performed single-cell RNA sequencing (scRNA-seq) of cells from mammary gland tissue isolated from PBS or IAV-primed female mice bred 21 days post-priming and analyzed at P5. Unsupervised clustering revealed 18 distinct cell populations across the epithelial (alveolar epithelial (AV) and myoepithelial cells), stromal (vascular endothelial and fibroblasts), neuronal, and immune mammary gland compartments (T cells, B cells, myeloid cells and distinct macrophages subsets; Fig. 5A, S10A). The overall cellular composition of the mammary gland was largely stable between IAV- and PBS-primed females, showing equivalent representation across most cell populations, with the exceptions within the immune compartment, with erythroid progenitor cells and inflammatory macrophages being significantly expanded in IAV-primed females (Fig. 5B). However, we noted transcriptional changes in many cell populations, suggesting that prior infection induces broad and long-lasting transcriptional remodeling of the mammary gland, despite no evidence of IAV infection of the mammary gland itself (Fig. S10B). Consistent with viral infections inducing robust IFNγ production, and with the broad responsiveness of diverse cell types to IFNγ ^54^, we find that maternal IAV priming establishes a lasting IFNγ-associated transcriptional signature across different cell populations within the mammary gland. This is especially evident in immune compartments including B cells, T cells, macrophages and endothelial cells populations, all of which show elevated expression of IFNγ-responsive genes (Fig. 5C, S10C). This suggests either sustained IFNγ production or durable imprinting of genes induced by IFNγ signaling during the original antiviral response. In contrast to this sustained interferon signature, NFkB associated signaling was reduced across multiple cell populations, suggesting that prior infection does not induce a generalized pro-inflammatory state, but instead establishes a selective, IFN-driven antiviral program that may preserve tissue homeostasis and enhance readiness to respond to viral challenge (Fig. 5D, S10D).

**Figure 5.**
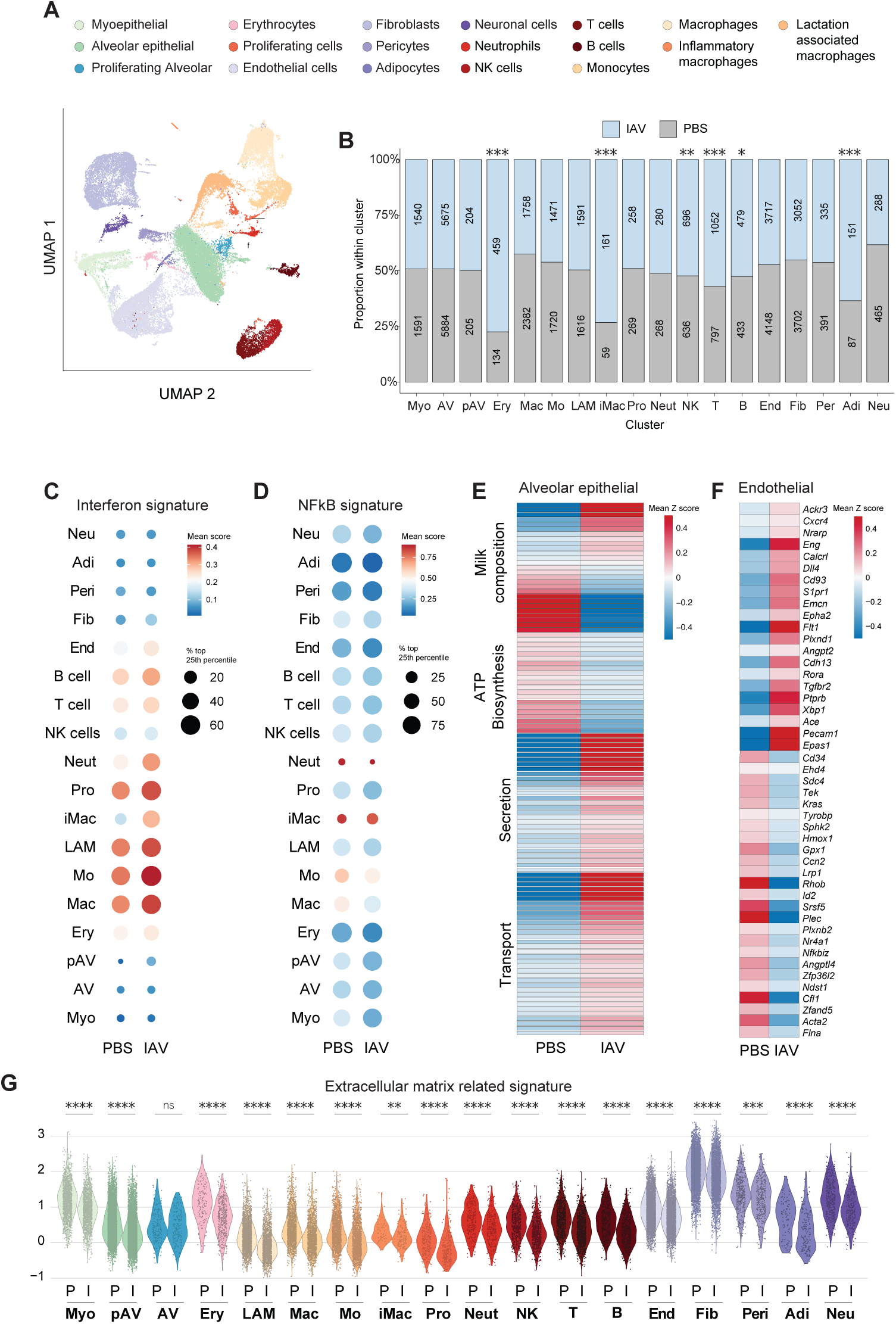
Maternal respiratory IAV exposure prior to pregnancy induces long-lived changes to the mammary gland. **(A-G)** Dams were infected with IAV (1 FFU, i.n.) or PBS and mated 21 days post infection. Cells from mammary glands of IAV and PBS-primed dams were analyzed by scRNA-seq at P5. **(A)** UMAP of clustered populations classified based on expression of canonical markers. Colors show indicated populations. PBS 28,638 cells; IAV 26,911 cells. **(B)** Proportion of cells within each cluster derived from each experimental group. Number inside bars indicates cell number of corresponding cluster per condition. **(C-D)** Differential gene expression analysis was performed on each cell cluster and scored for distinct modules (pathways). Dot Plot depicting per cluster average module scores for **(C)** interferon signaling and **(D)** NFkB signaling in indicated populations from mammary glands of IAV- or PBS-primed dams. **(E-F)** Heatmap of scaled expression of selected genes associated with **(E)** milk composition, ATP biosynthesis, secretion and transport in the alveolar epithelial cell cluster, and **(F)** vascular growth and function in the endothelial cell cluster. **(G)** Violin plot of selected extracellular matrix associated gene set across all clusters. Each dot represents one cell of indicated cluster. P= PBS, I= IAV. Cells for scRNA-seq analysis were isolated from mammary glands of 3 dams per condition. Differentially expressed genes with p adjusted value of <0.05 were used for analysis. See methods for scRNA-seq. In **(B) s**tatistical significance for differences in cell numbers per condition was determined using a hypergeometric test with FDR adjustment. In **(G)** statistical significance between conditions’ module score is determined using Kolmogorov-Smirnov tests. ns= not significant; *p<0.05, ** p<0.01, *** p<0.001.

Beyond immune associated changes, IAV exposure induced gene expression changes in stromal populations that suggested shifts in protein transport, energy consumption and tissue remodeling of mammary glands in IAV primed females. Indeed, the alveolar epithelial population showed the highest number of differentially expressed genes, with significant enrichment in pathways associated with protein secretion and transport, as well as changes in expression of genes associated with milk composition (Fig. 5E, S10E). We also saw decreased expression of genes involved in ATP biosynthesis (in particular, oxidative phosphorylation), as well as cholesterol biosynthesis suggesting a metabolic shift induced by IAV priming (Fig. 5E, S10E). Furthermore, we found the vascular population showed increased expression of genes involved in vascular remodeling, including upregulation of *Dll4, Nrarp, Angpt2, Plxnd1, Cxcr4* which are associated with tip-like sprouting angiogenesis phenotypes ^55–61^. These increases corresponded with the downregulation of *Tek, Ccn2, Acta2, Nr4a1*, which are involved in vascular stability ^62–65^ (Fig. 5F, S10F). Strikingly, nearly all clusters in the mammary glands of IAV-primed females displayed a coordinated downregulation of genes associated with extracellular matrix (ECM) production and remodeling (Fig. 5G). The ECM-related gene expression changes in nearly all populations in the mammary gland may reflect altered tissue remodeling dynamics that facilitate immune cell trafficking or antibody transport across the mammary epithelium. Alternatively, it could represent a shift in the balance between structural maintenance and secretory function that favors enhanced milk production, or a lasting consequence of inflammatory signaling, particularly by interferons, which have been associated with ECM gene alterations in other tissue contexts ^66–68^. This combination of transcriptional changes related to both vasculature and ECM suggests that the IAV primed mammary gland remodels to a state associated with increased vascularity that can facilitate permeability and increase access of maternal circulation to the mammary gland environment. Consistent with this notion, we find that immature erythrocytes, which are mostly present in circulation, were enriched in mammary gland from female mice that had been IAV primed compared to females primed with PBS cluster (0.54% in PBS vs 1.98% in IAV) (Fig. 5B). Overall, these changes indicate that prior maternal IAV exposure induces coordinated and long-lasting remodeling of the mammary gland to promote a tissue state optimized for the transfer of factors from circulation to milk for neonatal protection.

### The maternal-neonatal dyad confers intergenerational protection

Following birth, the enhanced antiviral immune state seen in late pregnancy is likely maintained and functionally extended into the perinatal period, a critical window in which maternal immunity continues to support neonatal defense. When neonates are highly susceptible to infection, pathogen exposure in the infant could simultaneously boost maternal immunity through re-exposure, in turn enhancing the transfer of protective factors back to the offspring through nursing, inducing a feedforward loop in which neonatal vulnerability is counterbalanced by amplified maternal protection. Such bidirectional immune communication could serve to rapidly amplify protective immunity within the maternal-neonatal dyad, ensuring survival during this vulnerable period. In line with this model, while IAV does not transmit readily between adult mice even by direct contact ^69^, we and others find that infected neonates are highly contagious and can transmit IAV^70^. Specifically, we detected viral transcripts in the lungs of dams exposed to infected neonates, but not in the mammary gland (Fig. 6A-B), indicating that neonatal-to-maternal transmission occurs via the respiratory route. Moreover, natural maternal exposure to IAV via the neonate was sufficient to confer protection to subsequent litters against lethal IAV (Fig. 6C-E). These results suggest that the bidirectional exchange of both virus and immunity within the maternal-neonatal dyad not only maximizes protection during the vulnerable period of early life, but could also result in an evolutionarily advantageous strategy for promoting immunity across multiple generation of offspring from the same parent.

**Figure 6.**
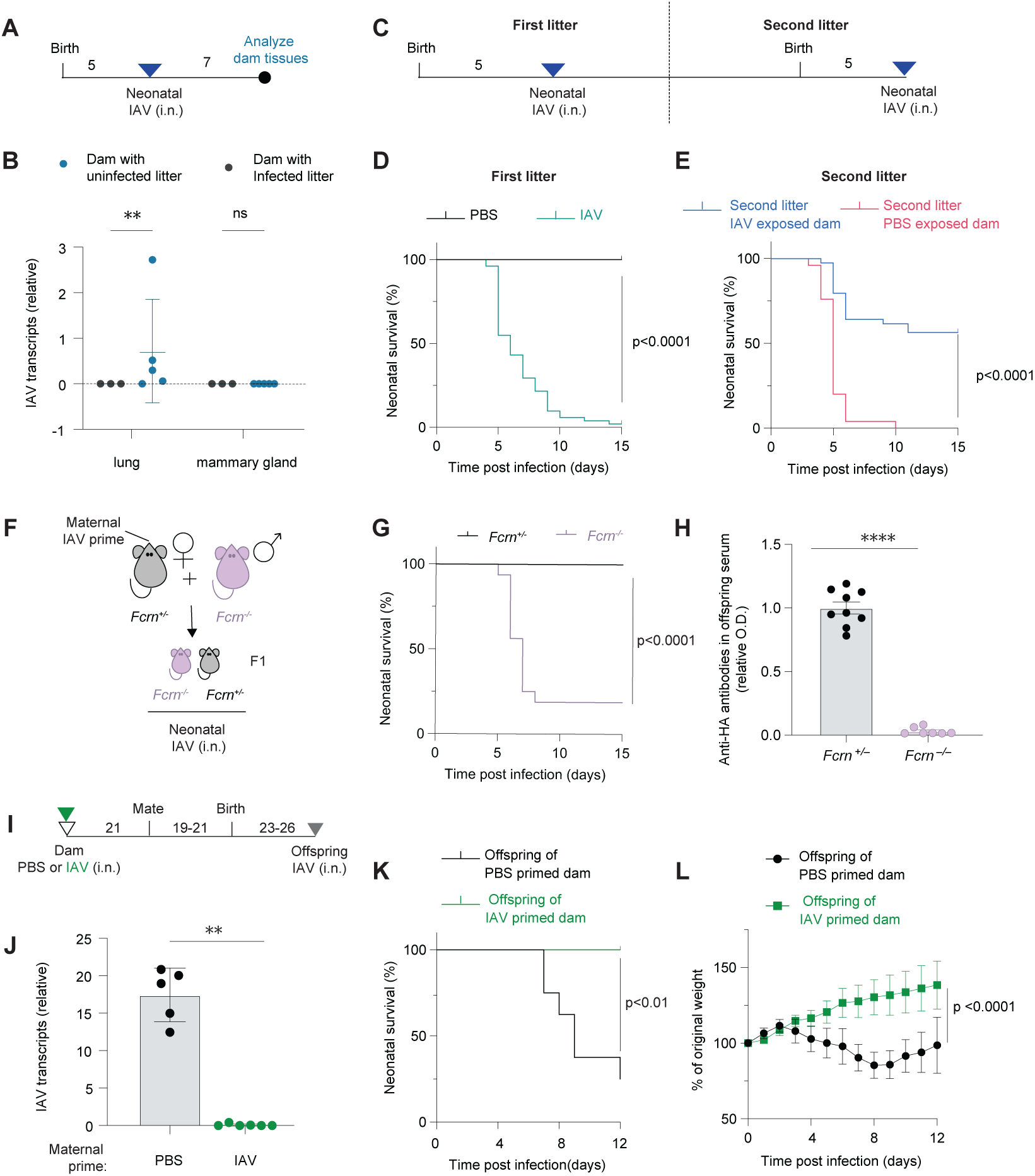
Maternal neonatal dyad establishes intergenerational long term antiviral immunity. **(A-B)** Neonates from naïve dams were infected with IAV (2FFU, i.n.) or PBS at P5. 7 days post infection dam tissues were analyzed for IAV viral transcripts. **(A)** Schematic of experimental setup. **(B)** IAV viral RNA transcripts in lungs and mammary glands quantified by RT-qPCR relative to lung *Actb* transcripts. Non-detectable values were assigned a zero. **(C-E)** Neonates from naïve dams (first litter) were infected with IAV (2FFU, i.n.) or PBS at P5. Dams were subsequently mated and neonates from the following litter (second litter) were infected with IAV (2FFU, i.n.) at P5. **(C)** Schematic of experimental setup. **(D)** Survival of first-litter neonates from naive dams following neonatal IAV infection of PBS at P5. PBS n=23; IAV n=51. **(E)** Survival of second-litter neonates from neonatal-IAV-exposed dam or neonatal-PBS-exposed dam, following neonatal IAV infection at P5. Neonatal-IAV-exposed dam n=39; neonatal-PBS-exposed dam n=25. **(F-H)** Heterozygous *Fcrn^+/-^* dams were infected with IAV (1FFU, i.n.) and mated 21 days post infection to homozygous *Fcrn^-/-^* males to generate *Fcrn^+/-^* and *Fcrn^-/-^* progeny. **(F)** *Fcrn*-deficient (*Fcrn ^-/-^*) and *Fcrn*-suficent littermate controls (*Fcrn^+/-^*) were infected (2 FFU, IAV i.n.). **(G)** Survival of *Fcrn ^-/-^* and *Fcrn^+/-^* following neonatal IAV infection at P5. *Fcrn^+/-^* n=22; *Fcrn^-/-^* n=16. **(H)** Serum from *Fcrn ^-/-^* and *Fcrn^+/-^* littermate controls were analyzed for HA-specific IgG antibodies by ELISA on P5. Optical density at 450 nm (O.D.) was normalized to the average O.D. of control group. Non-detectable values were assigned a zero. **(I-L)** Dams were infected with IAV (1 FFU, i.n.) or PBS and mated 18-21 days post infection. Juvenile offspring of IAV or PBS primed dams were infected with IAV (10 FFU, IAV i.n.) after weaning at 3 weeks old (P23-26). **(I)** Schematic of experimental setup. **(J)** IAV viral RNA transcripts in lungs infected offsprings quantified by RT-qPCR relative to lung *Actb* transcripts. Non-detectable values were assigned a zero. **(K)** Percent survival and **(L)** body weight change following juvenile IAV infection. Offspring of PBS-primed dam n =8; offspring of IAV-primed dam n=9. Dots represent data from individual mice, Bars show mean, error bars show SD **(B, J)** or SEM **(H)**. In **(L)** dot shows mean with error bars show SD. Statistical significance determined by 2-way ANOVA with Sidak’s multiple comparisons test **(B**), two-tailed unpaired t-test with Welch’s correction **(H, J),** survival Logrank test **(D, E, G, K)**, or 2-way ANOVA with fixed effects (type III) test, accounting time and experimental condition. **(L)**. ns= not significant; ** p<0.01, **** p<0.0001.

### Active neonatal transfer of maternal antibodies provides long-term antiviral immunity into early adulthood

Given the bidirectional exchange of virus and immunity within the maternal-infant dyad, and the central role of maternally derived IgG in mediating neonatal protection, we sought to define the mechanism by which these antibodies confer protection in the neonate. Specifically, we asked whether protection results from passive coating of neonatal mucosal surfaces by milk-derived IgG, or from active uptake of maternal IgG across neonatal epithelia into the circulation. Neonatal Fc receptor (FcRn) is shown to facilitate milk IgG transport across neonatal intestinal epithelium into circulation^71^. To directly test whether active uptake of milk IgG via FcRn is needed for neonatal protection, we primed *Fcrn^+/–^* dams which were then mated with *Fcrn^-/-^*males, generating both FcRn-sufficient (*Fcrn^+/–^*) and FcRn-deficient (*Fcrn^-/-^*) pups within the same maternal and nursing environment. At P5, pups were challenged with lethal IAV infection. We found that among *Fcrn^-/-^* neonates from primed dams succumbed to IAV infection while 100% of *Fcrn^+/–^* littermates survived through weaning (Fig. 6F-G). Aligned with the survival data, *Fcrn^-/-^* neonates lacked anti-HA IgG in serum (Fig. 6H), demonstrating that neonatal FcRn is essential for transporting milk IgG into neonatal circulation. This indicates that protection is mediated by systemic acquisition of maternal IgG rather than transient mucosal coating.

As we find that maternal IgG in milk is actively transported into neonatal circulation via the neonatal Fc receptor (Fig. 6H), this raises the possibility that maternally derived antibodies may persist and confer protection beyond the breastfeeding period itself. To test this, we asked whether neonates born to IAV primed dams retain protective immunity after weaning. We intranasally challenged offspring from PBS or IAV primed mice at postnatal day 23-26 with IAV (10FFU) and monitored viral burden and disease progression (Fig. 6I). Offspring from IAV primed dams were completely protected, displaying minimal to undetectable IAV transcripts in the lung 3 days post infection, compared to high viral burden in offspring from PBS-primed mice (Fig. 6J). Furthermore, offspring from IAV primed mice maintained continuous weight gain following normal growth trajectories following IAV infection (Fig. 6K-L). In contrast, offspring from PBS primed dams exhibited severe disease characterized by stunted growth, with the vast majority succumbing to infection within two weeks (Fig. 6K-L). This protection extended into early adulthood, with offspring showing significantly less weight loss at 8 weeks of age compared to those born to PBS-primed dams (Fig. S11A-C). These findings demonstrate that maternal immunity provides long-lasting defense against reparatory viral infection that persists into early life.

## DISCUSSION

Influenza viruses remain a major global health burden, particularly for young children who experience higher rates of hospitalization and complications ^72–76^. Current influenza vaccines are primarily inactivated formulations delivered intramuscularly, designed to elicit antibody responses predominantly against the highly variable head domain of the viral entry protein hemagglutinin (HA) of circulating viral strains (H1N1, H3N2 and B/Victoria lineages)^77,78^. While vaccines remain one of the most effective tools for reducing disease, the responses current formulations elicit can often be short-lived and vulnerable to antigenic drift, necessitating frequent vaccine updates ^79^. Additional strategies to improve vaccine efficacy have focused on the induction of mucosal immunity at the primary site of infection which may provide more effective and durable protection against respiratory viruses by enhancing local immune memory and antibody responses in the respiratory tract ^27,80,81^. In this context, our findings have important implications for understanding how long-lived immune responses established prior to pregnancy can be leveraged for neonatal protection. We demonstrate that preconceptual intranasal influenza infection allows for the transfer of protection to neonates, whereas priming via the footpad fails to protect offspring to subsequent influenza infection. This is likely due to higher influenza-specific antibody titers observed following intranasal exposure compared to footpad priming, suggesting the respiratory route is superior in generating protective humoral immunity. However, this may also reflect qualitative differences in immune responses generated at mucosal versus systemic sites, or simply the consequence of higher local antigen exposure from active viral replication in the respiratory tract. Regardless, our findings suggest that antiviral immune programs induced at mucosal sites may not only improve protection of the host but also enhance the transfer of protective immunity to offspring. Further, these results underscore the potential for promoting the use of mucosal vaccination or up-to-date intramuscular vaccination for reproductive-age individuals, given the apparent translatability as well as durability of antiviral immunity to neonatal protection.

The impact of prior respiratory viral infection on milk production or functional output has not been systematically examined in humans, and whether prior infection alters the quantity or secretory capacity of milk remains unclear, as direct measurements of milk production in the context of viral infection are limited and show minimal or no reduction ^82–84^. Much of what is known about viral infection-driven changes in milk production derives from agricultural models, where viral pathogens that directly infect the mammary gland and cause mastitis, like avian influenza virus (HPAIV) clade 2.3.4.4b H5N1, profoundly decrease milk yield. However, in contrast, systemic infection with Bovine Leukemia Virus, which infects B cells, is associated with increased milk production and changes in milk composition ^85,86^. Interestingly, we find that a prior viral infection in the lung induces a profound and unexpected changes in the mammary gland 47 days after infection that extends beyond direct immunological changes. In virus-primed mice, secretory luminal cells exhibit a shift away from ATP biosynthesis pathways toward programs supporting vesicular trafficking and secretory functions, suggesting an enhanced capacity for milk production and secretion. This was accompanied by stromal compartments showing downregulation of ECM components alongside signatures of active vascular remodeling and increased vascular permeability. Our analyses also revealed substantial changes of genes involved in local ECM remodeling. Although mechanisms governing plasma cell retention in the mammary gland remain poorly defined, studies in other tissues such as bone marrow and spleen have shown that plasma cell persistence and survival depend on integrin-mediated interactions with stromal compartments ^87–90^. Therefore, the downregulation of genes involved in ECM components in the mammary gland stroma may impair the retention and/or survival of IgA plasma cells, potentially explaining the reduction in total IgA in milk and decreased IgA plasma cells in the mammary glands of IAV-primed dams. Together, these coordinated changes point to a broader functional adaptation of the mammary gland, in which prior infection at a distal site both reshapes immune composition and enhances tissue programs that support the production and delivery of milk-borne factors. These findings add to a growing understanding that immune experiences actively shape mammary gland physiology beyond classical immune roles.

The epidemiological association between breastfeeding and reduced respiratory illness in young children is well established, including those caused by respiratory syncytial virus, SARS-CoV-2 and influenza virus ^91–94^. However, mechanistic understanding of how this protection is conferred has lagged, partly because of the focus on breastmilk’s most abundant antibody IgA ^36,95,96^. IgG, though present in breastmilk at lower concentrations and comparatively less stable at mucosal surfaces, is functionally active ^28,29,33,93^. Prior reports have documented virus specific IgG in breastmilk of previously infected or vaccinated individuals ^33–35,93,97^. In addition, virus-specific IgG has been shown to persist in breastmilk for up to eight months in those recovering from SARS-CoV-2 infection ^35^. Our results extend these observations by showing that IgG present in milk is functional and essential for conferring protection to neonates against respiratory infection in early life. Furthermore, structural analysis by EMPEM shows that milk IgG antibodies are enriched for binding HA’s stem esterase site. This suggests that the specific IgG populations present in milk can provide broad protection, as antibodies targeting these conserved regions have broadly neutralizing potential across antigenically distinct influenza strains ^98^. As cross fostering experiments demonstrated that milk is sufficient to protect offspring, even in the absence of *in utero* immune transfer, our study shows that IgG in milk is functionally active and capable of preventing viral infection in the neonate. This may be of particular importance in infants small for their gestational age or preterm birth. Transplacental IgG transfer occurs in the last trimester of gestation and peaks close to parturition, meaning that preterm infants are born with substantially lower maternally acquired IgG levels than their term counterparts ^99–102^. This indicates that breastfeeding, especially colostrum which is rich in antibodies ^103^, or oral IgG administration to neonates could be a therapeutic avenue for enhancing early-life immunity against respiratory viral pathogens for this particularly vulnerable population.

Based on our findings, active transport of IgG in infants through FcRn is necessary for its entry into neonatal circulation to allow protection. This is consistent with the established function of FcRn in mediating transcytosis of IgG across neonatal intestinal epithelium following ingestion of milk ^71^. Beyond this transport function, in adults FcRn contributes to effective humoral immunity throughout life by recycling IgG and extending its circulating half-life ^104,105^. As we find long term protection that persists beyond weaning, FcRn mediated salvage could also be involved in prolonging the effective half-life of maternally transferred IgG, well after the period of active milk consumption. Passively transferred maternal IgG could also shape the developing immune system in addition to providing immediate protection to neonates. Previous studies have shown maternal antibodies can alter neonatal follicular T helper dependent GC responses to vaccination and to the microbiota ^106,107^. In this context, the robust transfer of IgG observed in our study suggests that maternally derived antibodies, in those recovering from SARS-CoV-2 infection, have the potential of calibrating the magnitude and quality of neonatal immune responses with potential consequences for how offspring respond to infection and vaccination.

Lastly, our analyses show that preconceptual viral exposure reshapes the maternal B cell landscape during pregnancy. Compared to nulliparous controls, B cells from pregnant mice showed enhancement of antiviral programs, including an elevated interferon-primed state and increased germinal center activity, suggesting that during late pregnancy, B cell responses are amplified rather than suppressed. This is consistent with clinical observations that antibody-mediated autoimmune diseases, such as systemic lupus erythematosus, often flare during pregnancy ^108^. From an evolutionary perspective, this coupling of prior immune experience with pregnancy-associated immune remodeling likely serves to optimize protection of offspring against endemic pathogens encountered by the mother, providing sufficient neutralizing capacity to prevent infection upon early exposure and thereby delaying the initiation of adaptive immune responses until the neonatal immune system is more mature. Beyond providing protection, the transfer of maternal antibodies may also shape neonatal immune tolerance and curb responsiveness to these same endemic antigens ^107,109^. Thus, maternal immunological experience, pregnancy-associated immune adaptation, and tissue-level remodeling may converge to balance protection and immune education in early life. Our work establishes that preconceptual immune priming drives coordinated changes in maternal immunity and mammary gland remodeling to enhance offspring protection. These findings underscore the importance of considering reproductive-age individuals in the design of vaccination strategies and add to the growing mechanistic understanding on how maternal immunity shapes health and development of early life.

## Acknowledgments

We thank the Department of Animal Resources at Scripps Research for support with animal husbandry, as well as the Flow Cytometry Core and Genomics Core for technical assistance. We also thank all members of the Mendoza and Wiseman laboratories for technical input and discussion. This work is supported by the Prebys Foundation (A.M.) and National Institutes of Health (NIH) grants AG046495 (R.L.W) and AI136621 (A.B.W.). K.N.C. was supported by The Schimmel Family Endowed Fellowship Fund, and L.M.S. was supported by David C. Fairchild Endowed Fellowship, in the Skaggs Graduate School of Chemical and Biological Sciences. S.D. was supported by the Irvington Postdoctoral Fellowship from the Cancer Research Institute.

## Author contributions

K.N.C. and A.M. conceived the study. K.N.C., J.S.J., L.M.S., S.D., A.B.W., R.L.W. and A.M. designed and performed experiments with help from X.L. and N.W. K.N.C., J.S.J., L.M.S., S.D. and A.M. analyzed data and assembled the figures. A.B.W., R.L.W., and A.M. acquired funding. K.N.C., R.L.W. and A.M. wrote the paper. All authors reviewed and assisted in editing the paper.

## MATERIALS AND METHODS

### Mice

C57Bl/6J(Jax000664), *muMt^-^ (*Jax002288) *,Ighg^-^*(Jax038643) and *FcRn^-^* (Jax003982) were previously described ^28,104,110^ and were purchased from the Jackson Laboratory. *Tbx21^tdTomato-T2Acre^*mice were previously described ^111^. Generation and treatment on mice were approved by Scripps Research Institutional Animal Care and Use committee (protocol # 23-0004-1). All mice were bred and maintained at specific pathogen free (SPF) conditions, with controlled humidity and temperature, a 12h/12h light/dark cycle, with free access to water and standard chow diet. Female mice age 6 to 15 weeks old were used in all experiments. For timed mating experiments, males were introduced to female breeding cages at beginning of dark cycle and were removed the next morning. For cross-fostering experiments, pups were switched between experimental cages within 24h of birth. For survival experiments, mice were euthanized when moribund or when body weight decreases to more than 20% of original weight. For *FcRn^-^* mice experiments, neonates were identified by toe clipping and genotyped by PCR.

### Influenza infection

Influenza A/Puerto Rico/8/34 (IAV) was obtained from ATCC (#VR-95PQ). For adult intranasal infections, mice were anesthetized with 2.5% isofluorane and nasally inoculated with dosage described in text and figure legends of IAV diluted in 25uL phosphate buffered saline (PBS). For footpad priming experiments, mice were anesthetized with 2.5% isofluorane, and 50FFU (16,250 CEID50) IAV diluted in 25uL PBS was injected into hind limb footpad with 31-gauge needles. For neonatal intranasal infections, postnatal day 5 neonates were anesthetized with 3% isofluorane and nasally inoculated with 2FFU (650 CEID50) IAV diluted in 5uL PBS. Focus Forming Unit (FFU) for IAV stock was determined using focus forming assay on Madin-Darby Canine Kidney (MDCK) cells. Briefly, 25,000 MDCK cells were plated in 96 well flat bottom plates in D10 media [DMEM, 1%P/S, 10% FBS, 1% HEPES] and grown to confluency. Cells were washed with PBS to remove serum, and then inoculated with 10-fold serial dilutions of viral stocks in Flu Media [DMEM, 1mg/mL BSA, 1%P/S, 1ug/mL TPCK-trypsin]. After 1 hour incubation at 37°C, 100uL overlay medium [1:1 mixture of 2%methylcellulose and 2x MEM supplemented with 1%BSA, 2% P/S, 2% L-Glut, 2% HEPES, 2ug/mL TPCK-trypsin] was applied to each well. The plates were incubated for 24 hours at 37°C. After incubation, 100uL 4% paraformaldehyde was added to each well and incubated at room temperature for 20-30 minutes. Liquid was removed from each well, and the plate was washed with PBS 3-4 times. Viral foci were stained using a primary antibody against influenza A virus nucleoprotein (NP) (BioCell #BE0159, Clone HB-65) at 5ug/mL diluted in perm wash (PBS, 0.5g/L Saponin, 0.5g/L BSA), followed by HRP-conjugated secondary antibody (Southern Biotech 1010-05, 1:5000). After 1 hour incubation at room temperature, the plate was washed with 0.05% Tween PBS 3-4 times. Foci were visualized using KPL True Blue substrate (Seracare #5500-0049), and the plate was imaged with CTL Immunospot plate reader. Viral titers were calculated as FFU/mL = (Average number of foci) × (Dilution factor) / (Volume of inoculum in mL). Original stock of IAV from ATCC (#VR-95PQ, 2.1 x 10^11^ CEID_50_/mL) was determined as 6.4 x 10^8^ FFU/mL.

### RNA isolation and RT-qPCR

Tissues were harvested and total RNA was isolated using Direct-zol Miniprep kit (Zymo, R2051) according to manufacturer’s protocol. cDNA was synthetized from 1000ng RNA using High-Capacity cDNA Reverse Transcription Kit (Applied Biosystems, 4368814). RT-qPCR was performed using Power SYBR Green PCR Master Mix (Applied Biosystems, 4367659). Data analyzed as fold change normalized to *Actb* expression level.

Primers were purchased from IDT. Sequences target NP region of Influenza A/Puerto Rico/8/34 are:

Forward primer Sequence (5’->3’): TCAAAGGGACGAAGGTGCTC

Reverse primer Sequence (5’->3’): GCCCAGTACCTGCTTCTCAG

Samples with multiple or unidentifiable melting curves were assigned not detected (ND) and plotted as zero in graphs. When performing statistical analyses and calculating fold changes, pseudo count was added to all values to account for zero denominator.

### Tissue processing for cell isolation

For spleen and lymph nodes, tissues were mechanically dissociated, and cell suspension were passed through 100um strainer in FACS staining buffer [FSB; 1× PBS, 2% fetal bovine serum (FBS), 1 mM EDTA, and 10 mM HEPES]. Bone marrow was collected from one tibia per mouse. The tibia was harvested, two ends of the bone were cut, and the bone was placed into a PCR tube with a 22G puncture hole containing 50uL FSB. This PCR tube was put into a 1.7mL centrifuge tube and bone marrow cells were collected by brief centrifugation (7-9 seconds of short spin). For mammary gland, the inguinal, abdominal, thoracic and cervical mammary gland tissues were harvested, lymph nodes were removed, and tissues were weighed. Mammary gland tissue was submerged in 25mL digestion buffer [1x RPMI 1640, 2% FBS, 10mM HEPES, 1%Penicillin-Streptomycin, 1% L-Glutamine, 360U/mL Collagenase Type1, deoxyribonuclease I (DNase I; 1U/mL)] in 50mL centrifuge tubes with three ¼ inch ceramic beads (MP Biomedicals, 116540424-CF). For Lung, all lobes were collected and submerged in 3mL digestion buffer in 5mL snap cap tubes with one ¼ inch ceramic bead. The tubes were agitated at 250RPM for 20mins (mammary gland) or 45 mins (lung) at 37 °C. Digested tissue was then passed through a 70 µm cell strainer, and cells were pelleted by centrifugation (800 × *g*, 3 min at 4°C). Lymphocytes were subsequently enriched by density gradient centrifugation in 40% Percoll (ThermoFisher, cat. no. 45-001-747) in wash medium [1x RPMI 1640, 2% FBS, 10mM HEPES, 1%Penicillin-Streptomycin, 1% L-Glutamine]. For spleen, mammary gland and lung samples, red blood cells were lysed with ACK lysis buffer [155 mM ammonium chloride, 10 mM potassium bicarbonate, 100 nM EDTA pH 7.2]. Cells were then washed and resuspended in wash medium, and kept on ice until all samples were prepared for downstream processing.

### Flow cytometry

Mice were injected with 1.5ug of fluorophore-conjugated CD45 antibody in 200uL of PBS retro-orbitally under brief isoflurane anesthesia three minutes before sacrifice. Cells were isolated according to previous method section. Single cell suspensions were stained in 96 well V bottom plates. All washes were performed with 200uL FSB and all centrifugations were performed at 900 x g, 3 min at 4°C). For Influenza NP specific staining (tetramers obtained from NIH Tetramer Core Facility), cells were stained in Wash media supplemented with 5% FBS and 50uM beta-mercaptoethanol (Gibco) for 30mins at room temperature. Cells were then washed, and resuspended in 50uL/well Zombie NIR Fixable Viability dye with CD16/32 Fc block (BioLegend 101302; 1:200) diluted in 1 x PBS for 10 mins at 4°C. Cells were washed and proceeded with cell surface marker staining resuspended in 50uL/well FSB. For intracellular staining, CytoFix/CytoPerm kit (BD Bioscience; 554714) was used according to manufacturer protocol. For plasma cell nuclear staining of Blimp1, Foxp3/Transcription Factor Staining Buffer Set was used (eBioscience; 00-5523-00) according to manufacturer protocol. Before analysis, all samples were resuspended in 200uL FSB and passed through 100uM nylon mesh. All samples were acquired on 5 laser Cytek Aurora cytometer and analyzed using FlowJo (10.10.0).

### Antibodies

The following antibodies were used for flow cytometry experiments: CD16/32 (93), CD45R (RA3-6B2), CD90.2(53-2.1), CD62L(MEL-14), CD4(RM4-5), TCRγδ (GL3), CD45 (30-F11), TCRβ (H57-597), CD44 (IM7), IgD (11-26c.2a), CD11b(M1/70), CD8a (53-6.7), GL7 (GL7), CD64 (X54-5/7.1), CD95 (Jo2), CD38 (90), CD45 (30-F11), CD19 (6D5), IgA (C10-1), IgG2b (R12-3), IgM (II/41), IgG1 (X56), IgG2c (1077-30, Southern Biotech), Blimp1 (5E7), H2-Db | Influenza A NP 366-374 | ASNENMETM (NIH Tetramer Core), I-Ab | Influenza A NP 311-325 | QVYSLIRPNENPAHK (NIH Tetramer Core). The following antibodies were used for enzyme-linked immunosorbent assays: Anti-His-Tag-UNLB (SB194b), 6x-His Tag Recombinant Rabbit Monoclonal Antibody (RM146), Goat anti-Mouse Ig, Human ads-UNLB (1010-01, Southern Biotech), IgG1-HRP (1070-05, Southern Biotech), IgG2b-HRP (1090-05, Southern Biotech), IgG2c-HRP (1079-05, Southern Biotech), IgM-HRP (1021-05), IgA-HRP (1040-05, Southern Biotech), IgG3-HRP (1100-05, Southern Biotech), IgG-HRP (1030-05, Southern Biotech), Mouse IgM-UNLB (5300-01, Southern Biotech), Mouse IgG1-UNLB (5300-01, Southern Biotech), Mouse IgG2b-UNLB (5300-01, Southern Biotech), Mouse IgG2c-UNLB (0122-01, Southern Biotech), Mouse IgG3-UNLB(5300-01, Southern Biotech), Mouse IgA-UNLB(5300-01, Southern Biotech), Mouse IgG-UNLB (0107-01, Southern Biotech).

### Serum transfer

Immune serum collection: female nulliparous WT mice were inoculated with 10FFU (3250 CEID50) IAV (or PBS control) and serum was collected 21 days post infection. *μMT^-/-^* lactating dams were anesthetized with 2.5% isofluorane, and 200uL of immune or control naïve serum was injected retro-orbitally. For experiment testing antibody transfer efficiency from serum to milk, one dose of serum transfer was performed at postnatal day 1 or 2, and nursing dams were sacrificed 8-hour post transfer to harvest serum. Pups were sacrificed to collect breastmilk from stomach. For neonatal protection experiment, two doses of serum transfer were performed at postnatal day 3 and day 4, following neonatal lethal IAV infection at postnatal day 5.

### Enzyme-linked immunosorbent assay

Milk was harvested from pups’ stomachs at postnatal day 5 or 11. The milk paste was weighed and resuspend in 1x PBS to a concentration of 100 mg/mL. Samples were vigorously agitated by pipetting and vortexing, and then centrifuged at 10,000 x g for 5 min. Supernatants were collected and stored at −20°C until use. For HA-specific ELISA experiments, anti-His tag antibody was diluted in coating buffer [0.1M Sodium carbonate, pH 9.5 (8.4g NaHCO3, 3.56g Na2CO3 in 1L deionized water)] and coated onto high binding 96 well flat bottom microplate (Greiner 655061) at 50uL/ well overnight at 4°C. Plates were washed three times with ELISA wash buffer [PBS containing 0.05% Tween20] and then blocked with 150uL/well of 1% BSA in PBS for 2 hours at 37°C. Plates were then flicked and tapped dry on paper towels. Recombinant Influenza A H1N1 Hemagglutinin Protein (His-tag) was diluted in 0.5% BSA in PBS to a concentration of 0.25ug/mL, added to the plates 50uL/well and incubated for 1 hour at room temperature. After three washes with ELISA wash buffer, 50uL of breastmilk or sera (1:200 diluted in PBS) was added per well and incubated for at least 1 hour at 4°C. After three additional washes, HRP conjugated secondary antibodies diluted in ELISA wash buffer were added and incubated for 1 hour at 4°C. Plates were then washed five to six times with ELISA wash buffer, followed by incubation with TMB substrate. The reaction was stopped using 1M phosphoric acid. Absorbance was immediately read at 450nm optical density. Background obtained from the secondary antibody alone was subtracted. Absorbance readings less than background (with negative values) were considered not detected and assigned zero. To calculate fold change difference between control and experimental group, pseudo count of 0.01 was added to all values to account for zero denominator.

For Ig quantitative ELISA experiments, anti-mouse Ig antibodies were diluted in coating buffer to 1ug/mL and coated onto high binding 96 well flat bottom microplate at 50uL/well overnight at 4°C. Standards were prepared in PBS. Blocking and detection steps were performed as described above (serum dilution adjusted to 1:10,000 based on standard range 500ng/mL – 2.05ng/mL). Interpolation of unknown concentrations was performed using sigmoidal 4PL model in GraphPad Prism (Version 10.6.1).

### RNA-seq analysis

Mediastinal lymph nodes were isolated from mice (nulliparous vs pregnant E16.5-E17.5) 30-31 days after IAV priming. Three mice per condition were used, sequenced and analyzed. Naïve B cells (CD45^+^ Zombie NIR^-^ CD11b^-/lo^ CD64^-^ CD90.2^-^ CD19^+^ and/or B220^+^ IgD^+^ CD38^+^) and germinal center B cells (CD45^+^ Zombie NIR^-^ CD11b^-/lo^ CD64^-^ CD90.2^-^ CD19^+^ and/or B220^+^ IgD^-^CD38^-^ CD95^+^ GL7^+^) were obtained with fluorescence-activated cell sorting using a BD spectral sorter. 100,000-150,000 cells per population per mice were sorted into RLT Plus buffer supplemented with 2-mercaptoethanol (1:200, Sigma M3148). RNA was extracted from RNAeasy Plus Micro kit (Qiagen, 74134) according to manufacturer’s protocol. Library was prepared using NEBNext® Ultra™ II RNA Library Prep Kit for Illumina®, NEBNext® Multiplex Oligos for Illumina® (96 Unique Dual Index Primer Pairs Set 5). Sequencing was done by Scripps Research Genomic Core. 75 base pair paired-end reads (30 million per sample) were acquired on Element Aviti platform. RNA-seq reads were aligned to the reference mouse genome assembly mm39 using STAR RNAseq aligner^112,113^. Raw count of reads per gene was measured using R (4.3.3) and DEseq2 packages. BCR/TCR segment genes, as well as genes with zero counts in majority of samples are filtered out. For differential gene expression analysis, a cutoff of 0.1 was used to identify differentially expressed genes for each comparison. Hierarchical clustering and k-means method (k center=7) is used for clustering analysis.

### Electron Microscopy-Based Polyclonal Epitope Mapping

Electron microscopy-based polyclonal epitope mapping (EMPEM) as previously described ^52,53^. Serum and breastmilk samples were collected from PBS control and PR8-immunized groups of mice. Samples were heat-inactivated at 56°C for 1 hour prior to processing. Total IgG was isolated from serum and breastmilk using CaptureSelect IgG-Fc Affinity Matrix (Thermo Fisher Scientific). Bound IgG was eluted and subsequently digested into Fabs using activated papain in digestion buffer (100 mM Tris, 2 mM EDTA, 10 mM L-cysteine) for 4-5 hours at 37°C. The reaction was quenched with iodoacetamide at a final concentration of 30 mM. Fabs were separated from undigested IgG and Fc by size exclusion chromatography (SEC) using a Superdex 200 Increase 10/300 column (Cytiva) and concentrated using 10 kDa molecular weight cut-off Amicon ultrafiltration units (Millipore). Purified polyclonal Fabs from serum and breastmilk were incubated with recombinant PR8 HA trimer at a molar excess of Fab to trimer to ensure saturation of available epitopes. Immune complexes were purified by SEC to remove unbound Fab, and fractions corresponding to HA–Fab complexes were collected and concentrated for electron microscopy. Purified immune complexes were applied to glow-discharged 400-mesh copper grids coated with carbon film and stained with 2% uranyl formate. Micrographs were collected on a Thermo Fisher Talos F200C 120 kV transmission electron microscope equipped with a Ceta camera and Thermo Scientific Smart EPU software^114^. Datasets of single particles were collected for each condition, where ∼100,000 particles were picked and subjected to iterative rounds of reference-free 2D classification to separate apo HA trimers from Fab-bound complexes. Classes showing Fab density were selected and subjected to 3D classification and refinement. Individual Fab densities were segmented from the 3D reconstructions, and composite polyclonal epitope maps were assembled by resampling each unique Fab class onto the HA trimer initial model to generate a complete epitope landscape for each sample. Data processing was performed using RELION 4.0 software package^115^ and UCSF ChimeraX^116^. Epitope specificities were assigned based on the location of Fab density relative to the HA trimer structure, with sites classified as head, side head, esterase, stem, or protomer–protomer interface, consistent with previously defined HA epitope nomenclature^51^.

### Single cell RNA seq analysis

Mammary gland tissues were isolated and dissociated into single-cell suspensions as described in previous section. Dead cells were removed using MojoSort Mouse Dead Cell Removal Kit (BioLegend, 480157) according to manufacturer’s protocol. Cells were then fixed, and library preparation was performed using Chromium Next GEM Single Cell Fixed RNA Sample Preparation Kit (10x Genomics PN-1000414) following manufacturer’s instructions. Cells from three mice per condition were pooled (750,000 cells/condition), barcoded and multiplexed prior to sequencing. Libraries were sequenced on an Element Aviti platform to generate 75-base pair paired-end reads, yielding approximately 1 billion total reads. After preprocessing, 28,638 cells from PBS samples and 26,911 cells from IAV samples were retained for analysis. Reads were aligned to the mm10-2020-A reference transcriptome and demultiplexed using Cell Ranger (10x Genomics, v9.0.1). Quality control filtering was performed by retaining cells/nuclei were with more than 200 detected genes, fewer than 20,000 total RNA molecules, and less than 15% mitochondrial transcript content. Downstream analyses were performed in R (v4.3.3) using Seurat pipeline (v5.2.1). Heterotypic doublets were identified and removed using DoubletFinder^117^. A K-nearest neighbor graph was generated based on the PCs using FindNeighbors function, and clusters were identified using the Louvain algorithm implemented in FindClusters. An initial clustering resolution of 0.1 was used for broad lineage identification, and the resolution was subsequently increased up to 1.5 to achieve finer cluster granularity. Conserved markers across conditions were identified using FindConservedMarkers. Cluster identities were manually annotated based on CellMarker2.0 ^118^, Annotation of cell types (ACT) ^119^, the SingleR package ^120^ with ImmGenData and MouseRNAseqData database and peer-reviewed literature ^17,18,121–124^. Differential expression (DE) analysis was performed using DElegate package, which assigns cells to pseudo-replicates followed by differential expression testing with DEseq2 method^125^. Genes expressed in fewer than 10% of cells within a given cluster were excluded from analysis. Genes with an adjusted p-value <0.05 were considered statistically significant. Gene Set Enrichment Analysis (GSEA)^126^ and Metascape^127^ were used to interpret gene expression data. Pathway-associated transcriptional programs (gene sets listed in Supplementary Table 1) were evaluated using Seurat’s *AddmoduleScore* function. For IFN*γ*-, Nf*κ*B-response signature, module scores were summarized within each cluster by calculating the mean and median score, as well as the proportion of cells with high signature scores, defined as values above the third quartile across all cells. Descriptive differences in ECM-module scores between conditions were assessed at single cell resolution using Wilcoxon rank-sum or Kolmogorov-Smirnov tests, with individual cells treated as observations. Cell proportion within each cluster in IAV and PBS condition were evaluated with hypergeometric test with FDR adjustment.

### Statistical analysis

All statistical analyses and data graphing, except RNA seq, were performed using GraphPad Prism (Version 10.6.1). Normal distribution was assumed for data distribution. Mice were randomly assigned to experiments and each experiment is age matched. For influenza specific enzyme-linked immunosorbent assays and viral genome copies RT-qPCR experiments, pseudo count was added to all values to account for zero denominator when calculating fold changes. *p ≤ 0.05;**p ≤0.01;***p ≤ 0.001, ****p ≤0.0001; ns, not significant.

## Data, code, and materials availability

Raw and processed sequencing data for bulk RNA-seq and scRNA-seq will be available through GEO. Three-dimensional maps and models for the EM analysis have been deposited to the Electron Microscopy Databank (EMDB) (http://www.emdatabank.org/) (EMDB IDs: EMD-77394, EMD-77395, EMD-77396, EMD-77398) All ns-EM EMDB accession numbers are listed in Supplementary Table 2. Any additional information required to reanalyze the data reported is available upon request.

---

**Table S1.**
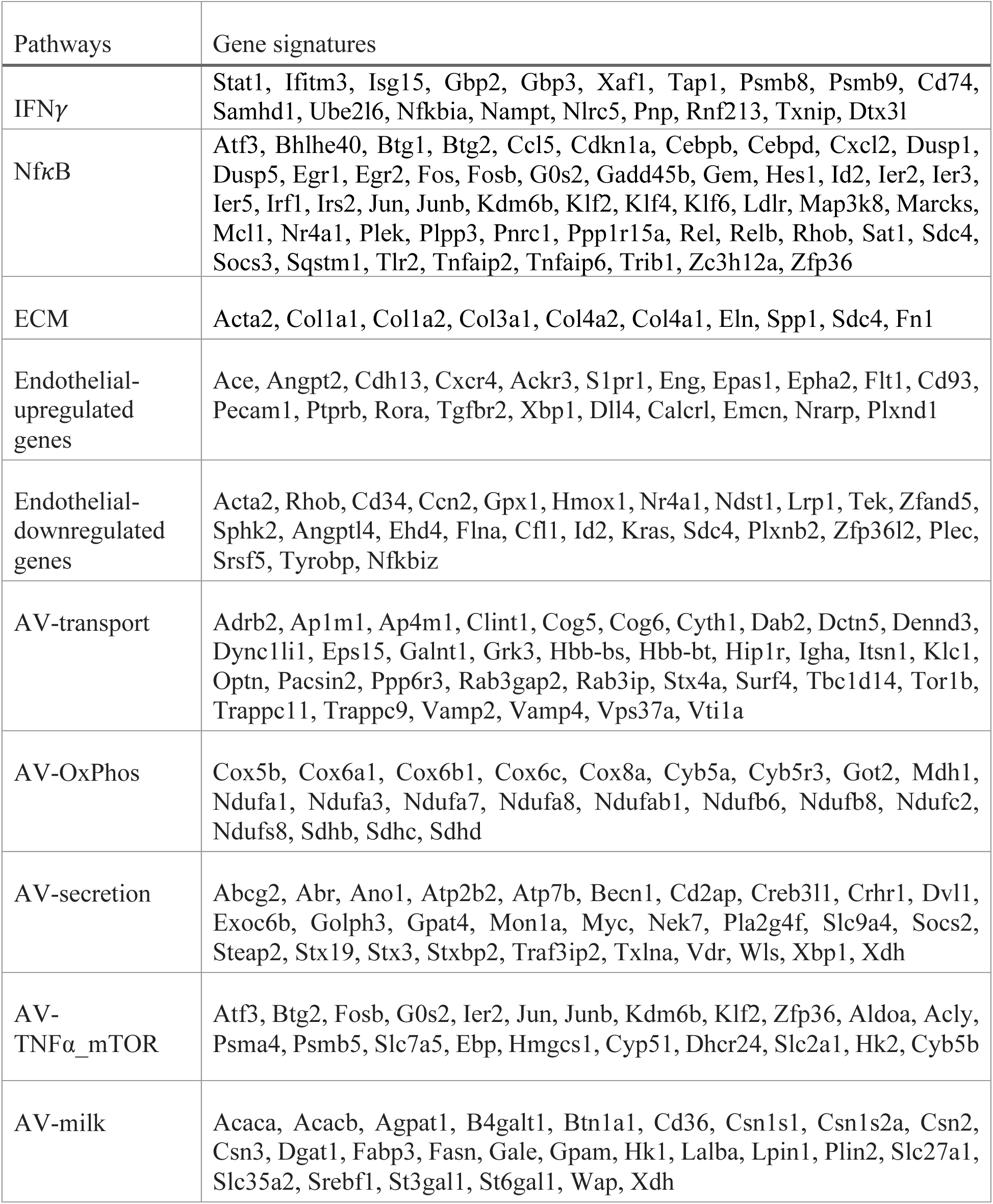
Pathway-associated transcriptional programs gene signatures used in single cell RNA sequencing analyses.

**Table S2.** Ns-EM Electron Microscopy Data Bank deposition information. Maps are accessible at emdataresource.org using the listed codes. Additional maps can be found on the “Download” tab of each entry.

| **EMDB Code** | **Influenza HA Strain** | **Sample ID** | **Map (C1 Symmetry)** | **Polyclonal Antibody Epitope** | **Contour** |
| --- | --- | --- | --- | --- | --- |
| EMD-77394 | A/Puerto Rico/08/1934 | PBS Serum Pooled | Main Map + Half Maps 1 & 2 | Stem | 0.02 |
| EMD-77395 | A/Puerto Rico/08/1934 | PBS Breastmilk Pooled | Main Map + Half Maps 1 & 2 | Stem | 0.02 |
| EMD-77396 | A/Puerto Rico/08/1934 | PR8 Serum Pooled | Main Map + Half Maps 1 & 2 | Stem | 0.02 |
|  |  |  | Additional Map | Esterase | 0.02 |
|  |  |  | Additional Map | Side Head | 0.02 |
|  |  |  | Additional Map | Head | 0.02 |
| EMD-77398 | A/Puerto Rico/08/1934 | PR8 Breastmilk Pooled | Main Map + Half Maps 1 & 2 | Esterase | 0.02 |
|  |  |  | Additional Map | Stem | 0.02 |

**Figure S1.**
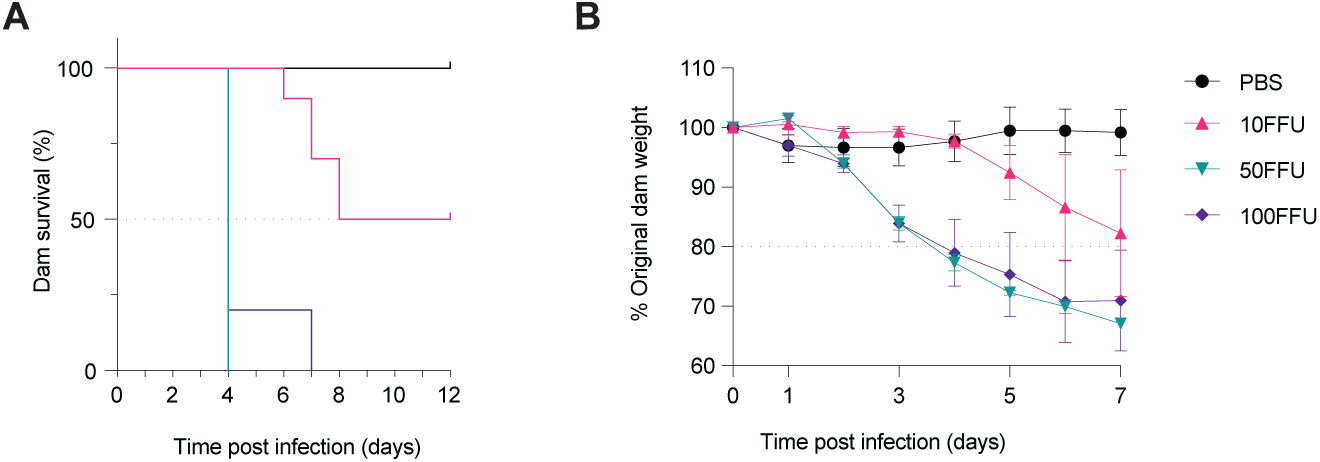
Dose response of PR8 IAV intranasal infection in adult mice. **(A)** Percent survival and **(B)** percent body weight loss of adult female mice following intranasal IAV of indicated dose or PBS. PBS n=6; 10FFU n=4; 50FFU n=3; 100FFU n=5. Representative of more than 2 experiments. Dots represent average data from each time point with SD **(B)**.

**Figure S2.**
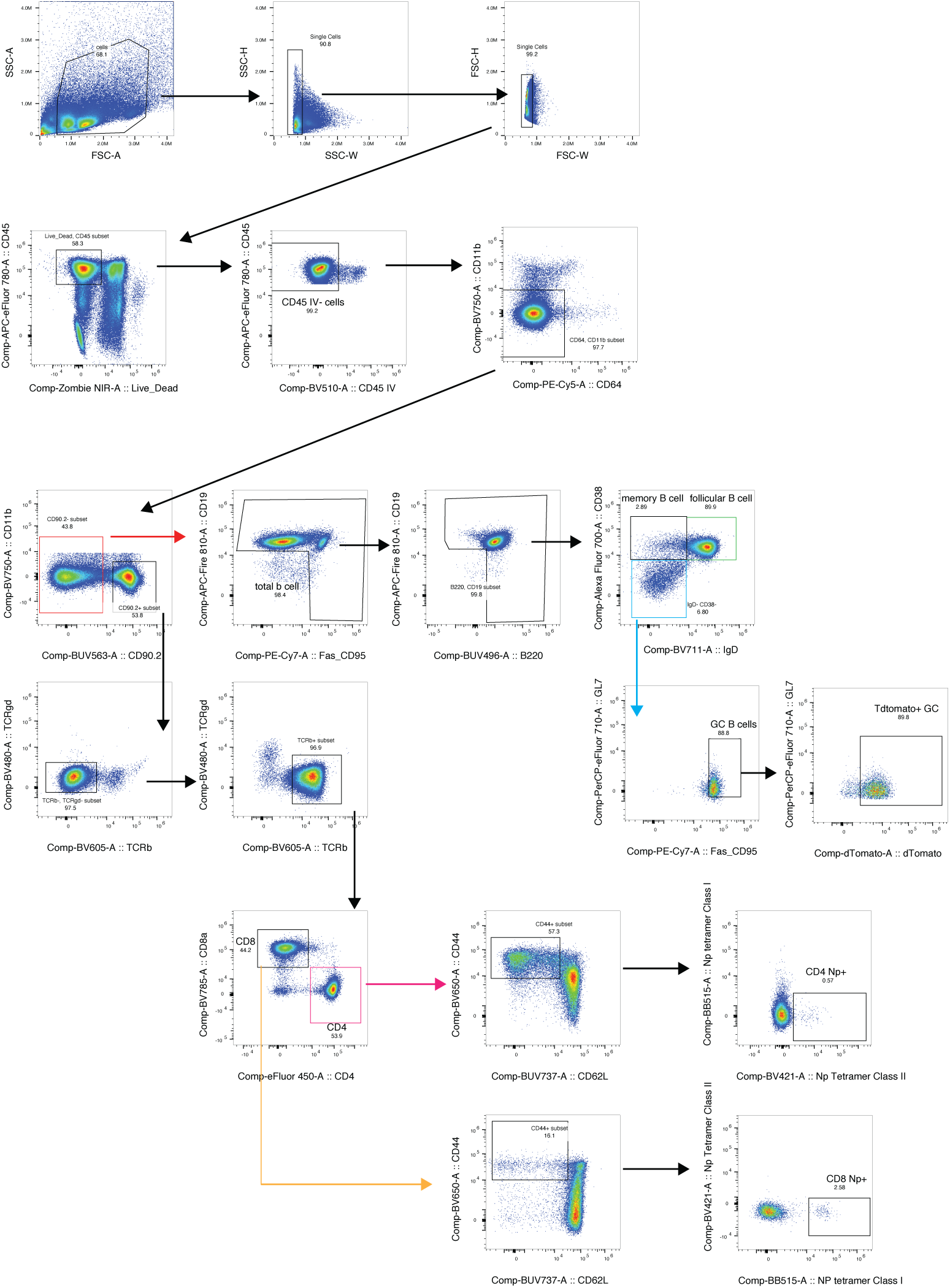
Gating strategy. Representative flow cytometry gating strategy for identifying immune cell populations, with NP tetramer used to identify antigen-specific T cells. In mice carrying *Tbx21^cre-TdTomato^*, TdTomato was used to examine T-bet expression.

**Figure S3.**
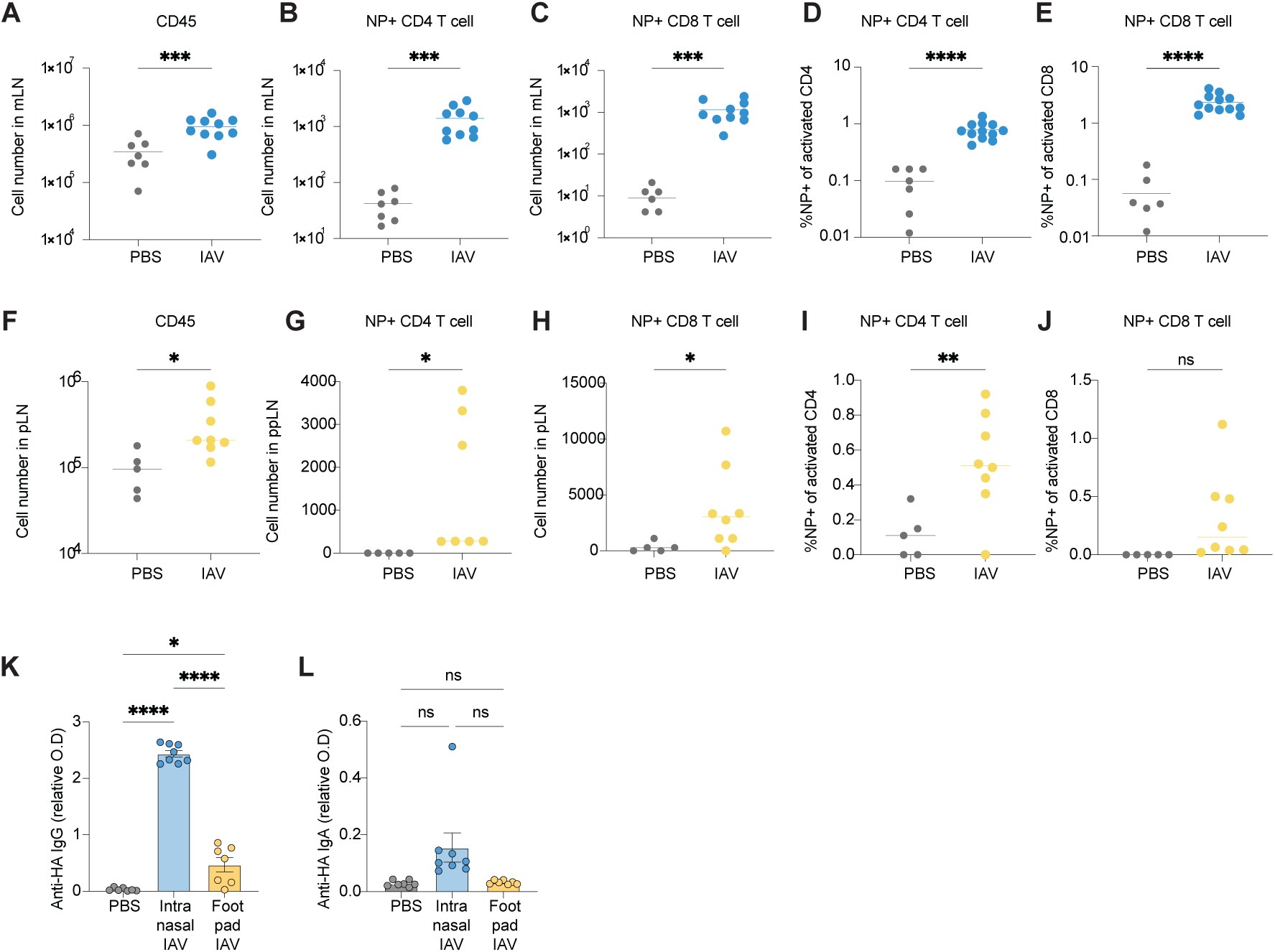
Route dependent induction of influenza specific maternal adaptive immunity. **(A-E)** Dams were infected with IAV (1 FFU, i.n.) or PBS and mated 21 days post infection. Mediastinal LNs were collected at P5 and analyzed by flow cytometry. **(A-C)** Cell numbers of **(A)** CD45+ cells, **(B)** influenza nucleoprotein (NP) specific CD4+, and **(C)** NP+ CD8+ cells. (**D-E**) Frequency of NP+ CD4+ of activated CD4 T cells **(D)** and and NP+CD8+ T cells **(E)** of activated T cells. **(F-J)** Dams mice were primed in the hind footpad with IAV (50 FFU) or PBS and analyzed 7 days post injection. Popliteal LN was collected at P5 and analyzed by flow cytometry. **(F-H)** Cell numbers of **(F)** CD45+ cells, **(G)** influenza nucleoprotein (NP) specific CD4+, and **(H)** NP+ CD8+ cells. **(I-J)** Frequency of NP+ CD4+ of activated CD4 T cells **(I)** and and NP+CD8+ T cells **(J)** of activated T cells. **(K-L)** Dams were inoculated with PBS (i.n.), intranasal IAV (1 FFU) or footpad IAV injection (50 FFU). Relative titers of HA-specific IgG **(K)** and IgA **(L)** antibodies level in sera 21 days post inoculation, quantified by ELISA. Optical density at 450 nm (O.D.) was normalized to the average O.D. of control group per isotype. Non-detectable values were assigned a zero. Dots represent data from individual mice **(A-L)**. Scatter dot plots show mean. Bar plots show mean, error bars show SEM (**A-L**). Statistical significance determined by two-tailed unpaired t-test with Welch’s correction **(A-J)**, or 2-way ANOVA with Sidak’s multiple comparisons test **(K, L)**. ns= not significant; *p<0.05, ** p<0.01, *** p<0.001, **** p<0.0001.

**Figure S4.**
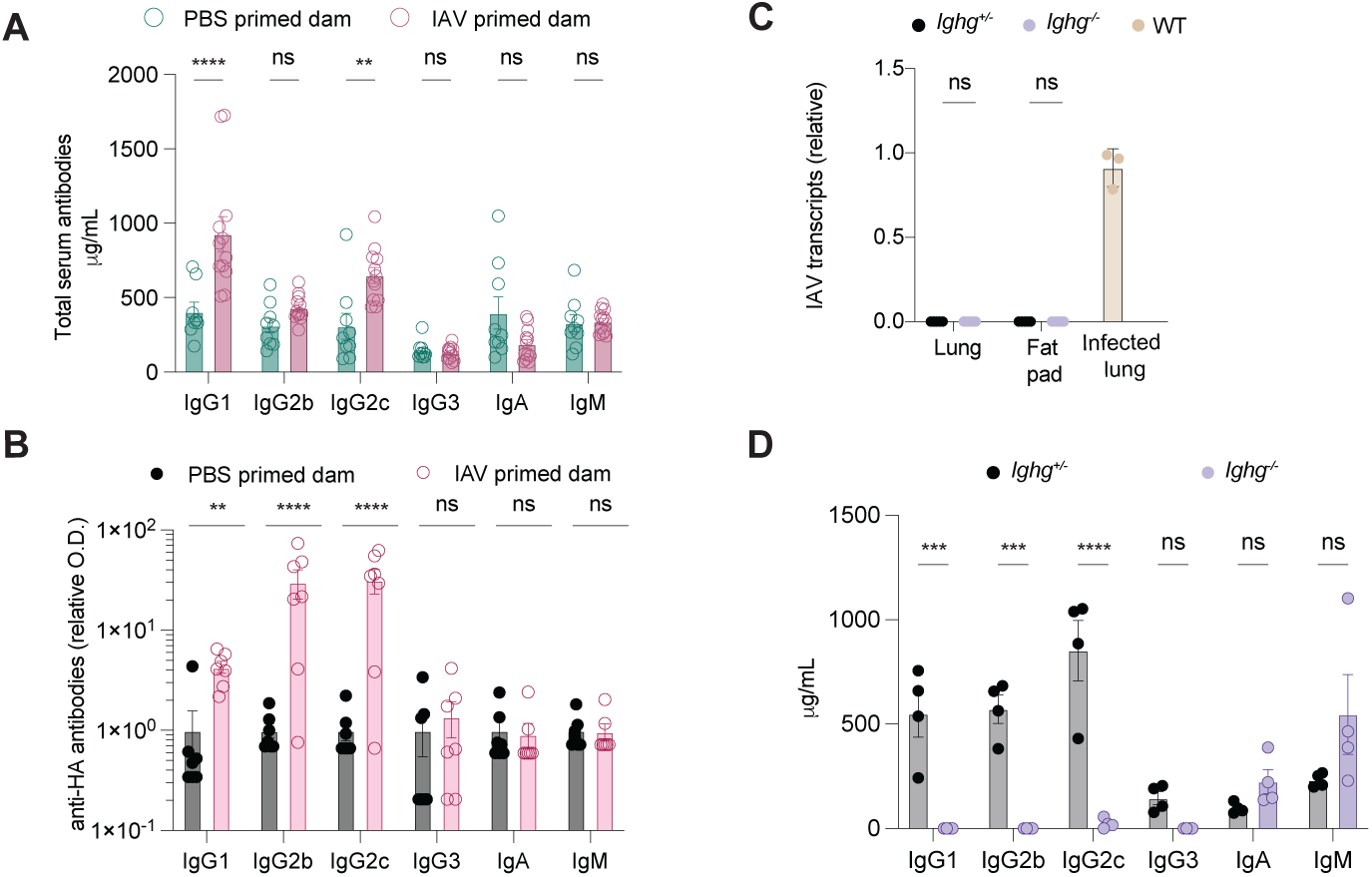
Maternal IAV-specific antibody response is predominantly IgG. **(A-B)** Dams were infected with IAV (1 FFU, i.n.) or PBS and mated 18-21 days post infection. **(A)** Concentration of total antibodies of indicated isotypes in sera of dams at P5, quantified by ELISA. **(B)** Relative titers of influenza hemagglutinin (HA) specific antibodies of indicated isotypes in milk quantified by ELISA on P11. Non-detectable values were assigned a zero. Optical density at 450 nm (O.D.) was normalized to the average O.D. of control group per isotype. **(C-D)** *Ighg^-/-^* and control (*Ighg^+/-^)* mice were infected with IAV (1 FFU, i.n.) and mated 21 days post infection. **(C)** IAV viral RNA transcripts in lungs and mammary fat pads of IAV infected (1FFU, i.n.) *Ighg^-/-^* and control (*Ighg^+/-^)* dams, quantified 14 days post infection by RT-qPCR relative to lung *Actb* transcripts. IAV infected (1FFU, i.n.) lungs of mice collected 3 days post infection were used as positive control for viral transcripts. Non-detectable values were assigned as zero. **(D)** Concentration of total antibodies of indicated isotypes in sera of dams at P5, quantified by ELISA. Dots represent data from individual mice, bars show means, error bars show SEM (**A, B, D)** or SD **(C)**. Statistical significance determined by 2-way ANOVA with Sidak’s multiple comparisons test (**A, B, D**). ns= not significant; ** p<0.01, *** p<0.001, **** p<0.0001.

**Figure S5.**
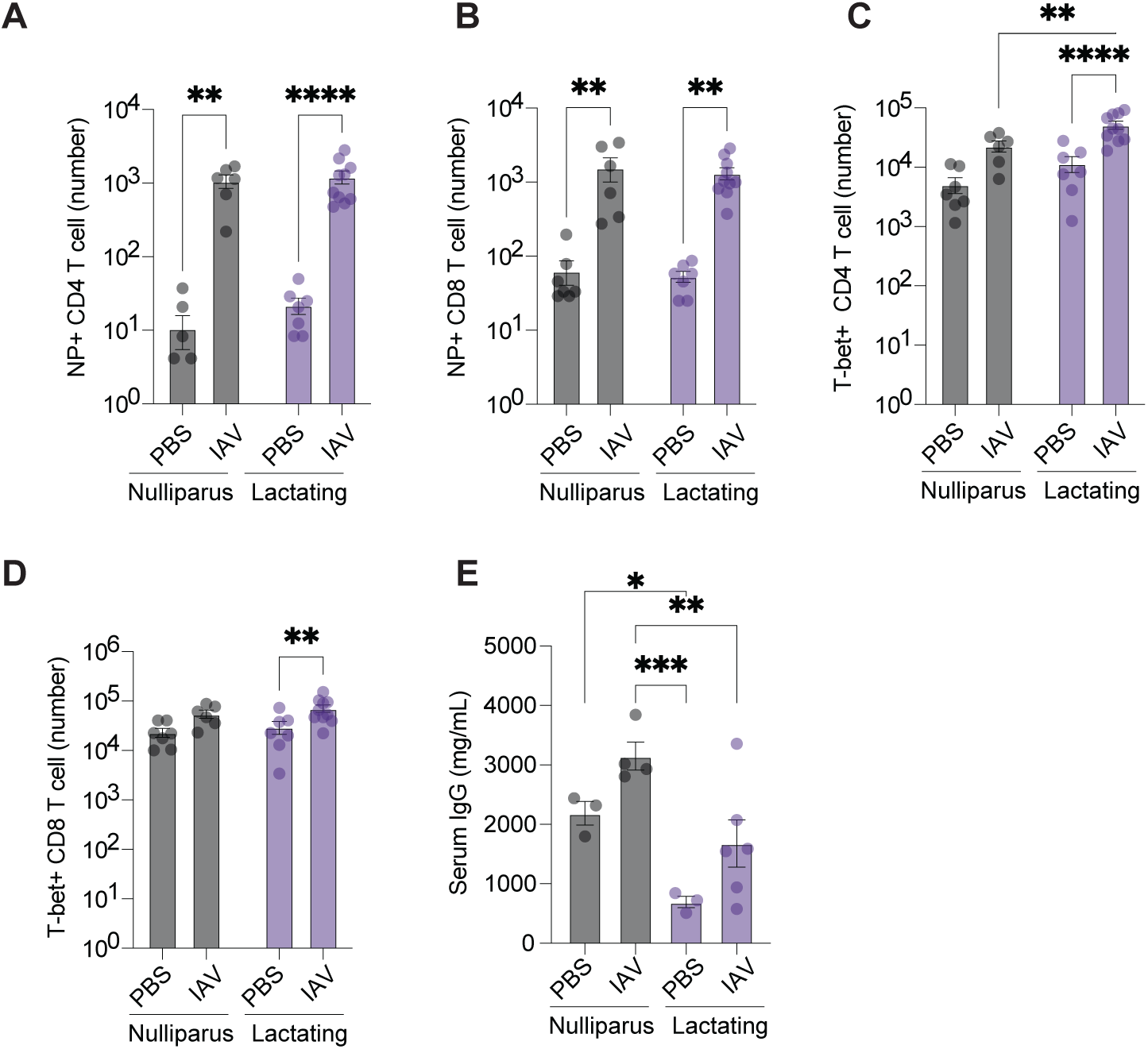
Lactation enhances antiviral, influenza specific T cell responses. **(A-E)** *Tbx21^TdTomato-Cre^* mice were infected with IAV (1 FFU, i.n.) or PBS and mated or left unmated 21 days post infection. T cells were analyzed by flow cytometry at P5 from medLN following CD45 I.V. labeling. Cell number of **(A)** NP+ CD4 T cells, **(B)** NP+ CD8 T cells, **(C)** Tbet+ CD4 T cells and **(D)** Tbet+ CD8 T cells. **(E)** Concentration of total IgG antibodies in sera of nulliparous and lactating dams at P5, quantified by ELISA. Dots represent data from individual mice, bars show means, error bars show SEM **(A-E)**. Statistical significance was determined by 2-way ANOVA with Sidak’s multiple comparisons test **(A-E)** or two-tailed unpaired t-test with Welch’s correction **(D)**. *p<0.05, ** p<0.01, *** p<0.001, **** p<0.0001.

**Figure S6.**
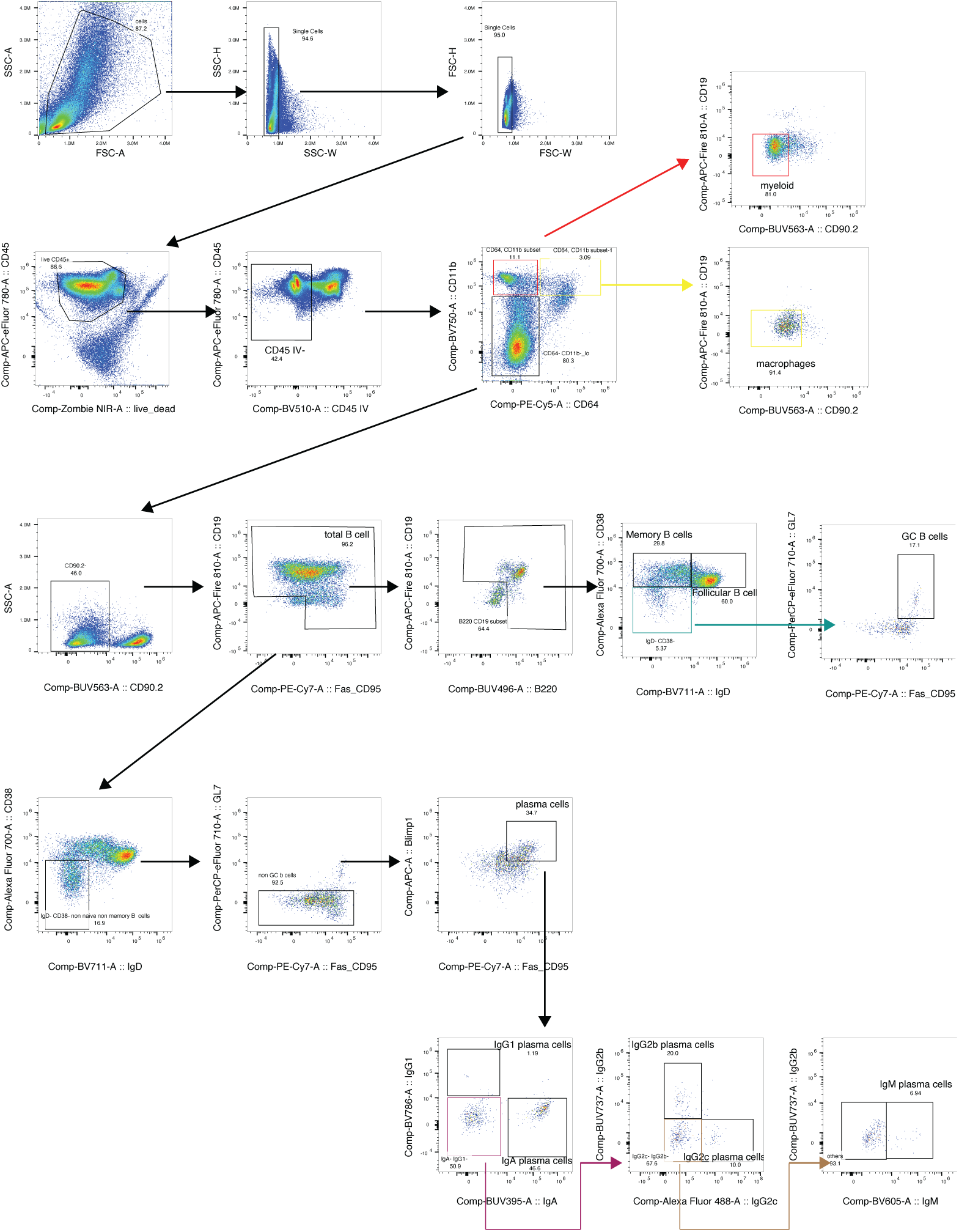
Gating strategy. Representative flow cytometry gating strategy for identifying myeloid cells and plasma cell populations.

**Figure S7.**
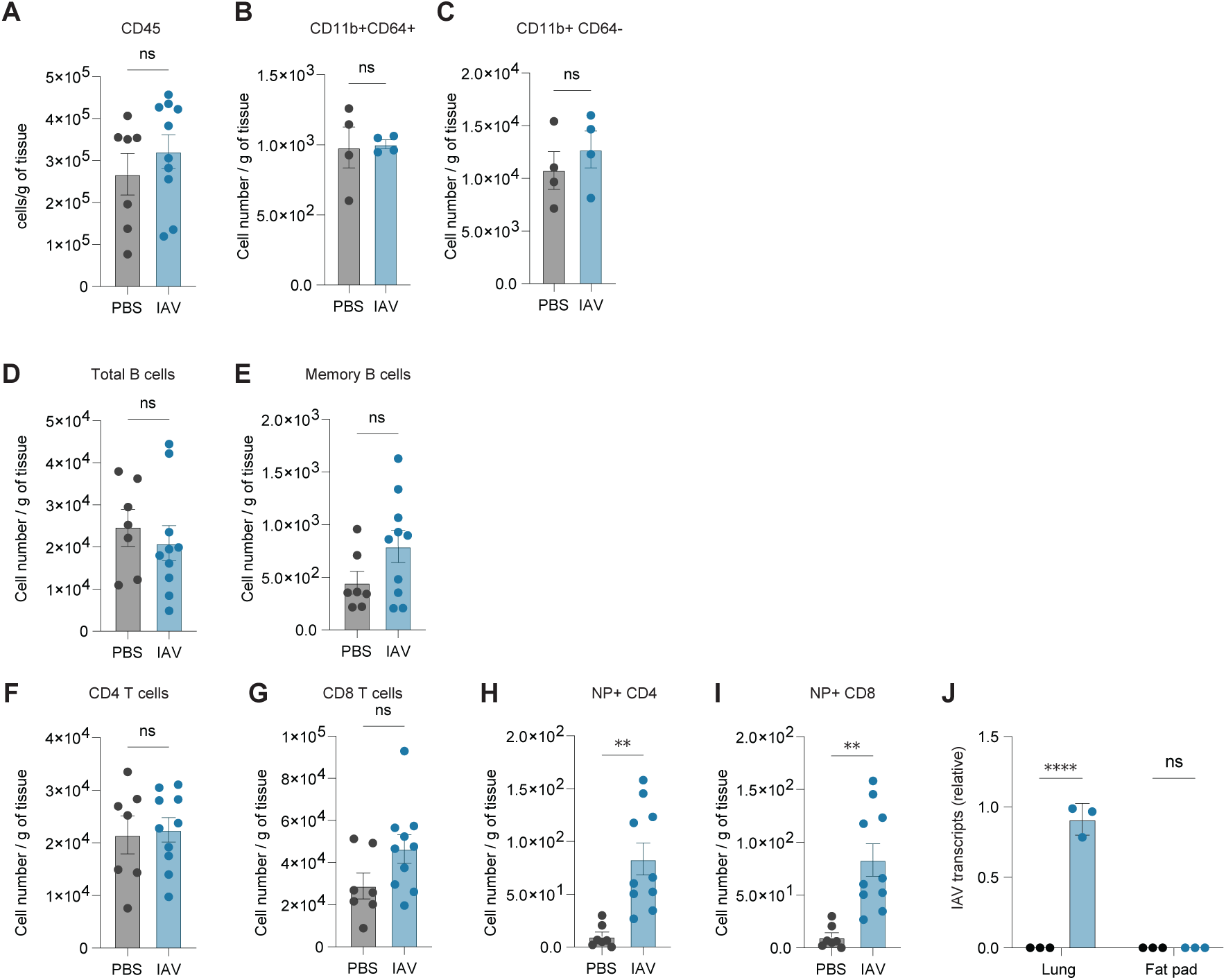
Preconceptual Respiratory Maternal IAV exposure induces long-term cellular changes in the mammary gland. **(A-I)** Dams were infected with IAV (1 FFU, i.n.) or PBS and mated 21 days post infection. Immune cells were quantified in mammary glands at P5 by flow cytometry following CD45 I.V. labeling. Numbers of nonvascular total CD45+ cells **(A)**, CD11b+ CD64+ cells **(B)**, CD11b+ CD64-cells **(C)**, total B cells **(D)**, memory B cells **(E)**, CD4+ T cells **(F)**, CD8+ T cells **(G)**, influenza nucleoprotein (NP) specific CD4+ T cells **(H)** and NP specific CD8+ T cells **(I)**. **(J)** Dams were infected with IAV (1 FFU, i.n.) or PBS. IAV viral RNA transcripts in lungs and mammary fat pads were quantified 3 days post infection by RT-qPCR relative to lung *Actb* transcripts. Non-detectable values were assigned a zero. Dots represent data from individual mice, bars show means, error bars show SEM (**A-I)** or SD **(J)**. Statistical significance determined by two-tailed unpaired t-test with Welch’s correction **(A-I),** or 2-way ANOVA with Sidak’s multiple comparisons test **(J)**. ns= not significant, ** p<0.01, **** p<0.0001.

**Figure S8.**
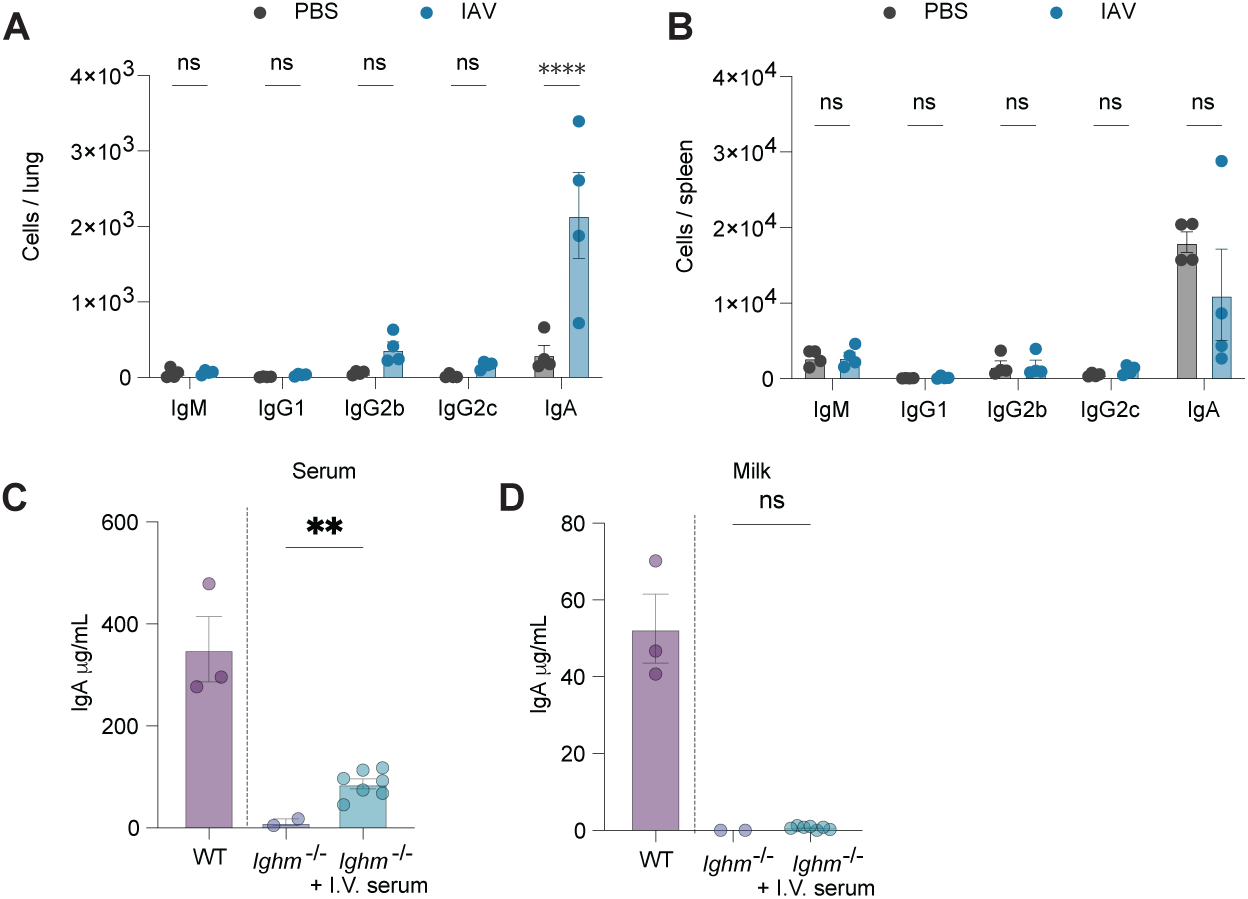
Preconceptual Respiratory Maternal IAV exposure induces systemic cellular changes in lactating dams. **(A-B)** Dams were infected with IAV (1 FFU, i.n.) or PBS and mated 21 days post infection. Plasma cells were analyzed by flow cytometry were quantified by flow cytometry at P5. Number of plasma cells of indicated antibody isotypes in lung (**A)** and spleen **(B)**. **(C-D)** Serum from WT mice was transferred to *Ighm⁻/⁻* dams. Eight hours after transfer, IgA antibody was quantified in serum **(C)** and milk **(D)** of *Ighm⁻/⁻* recipient dams by ELISA.

**Figure S9.**
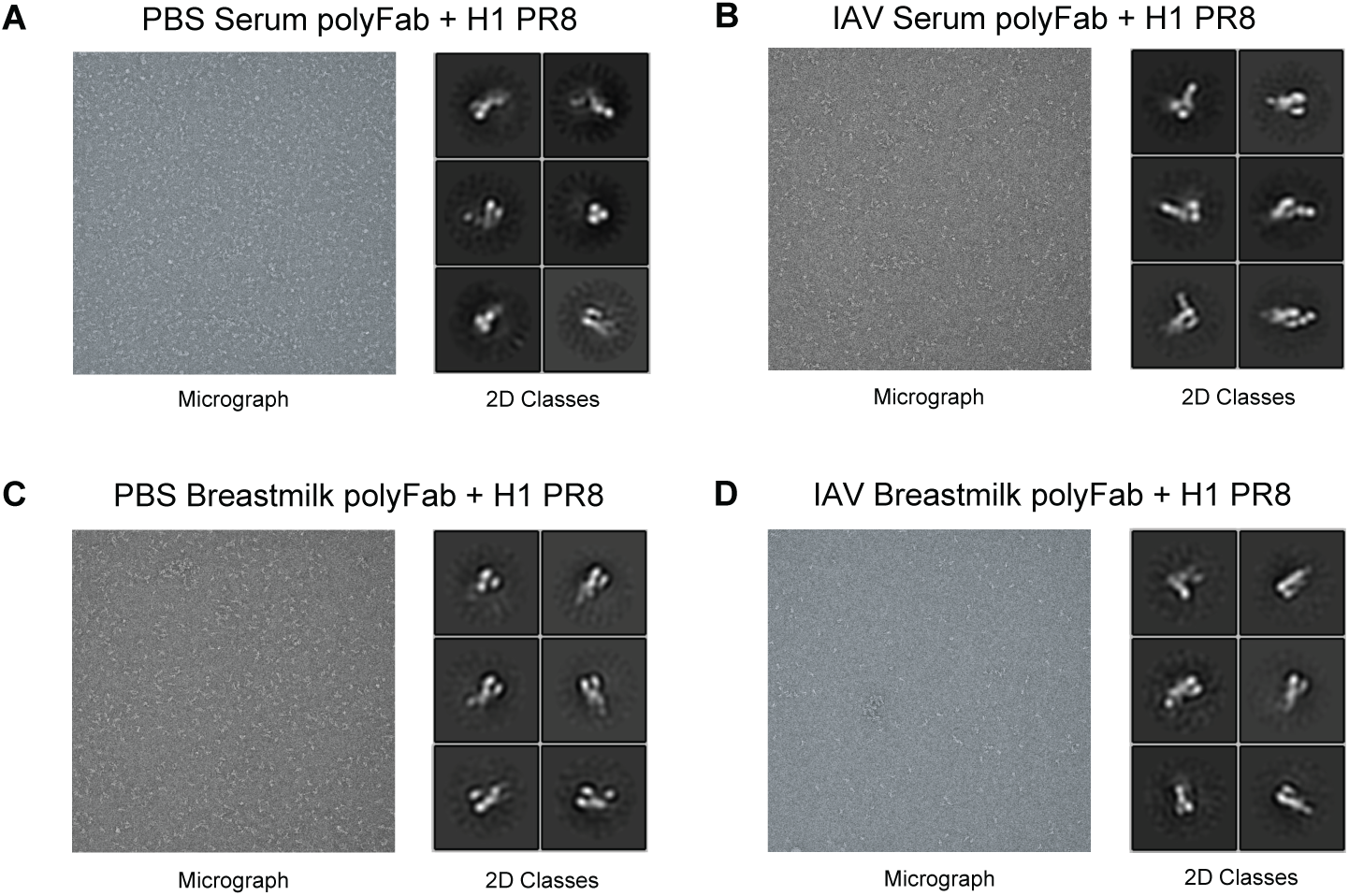
Extended ns-EM analysis for polyclonal immune complexes with H1 PR8 trimer. Representative micrographs and 2D class averages from the ns-EM datasets were used for the generation of composite figures presented in Figure 2I-J. **(A-B)** Sera from PBS- and IAV-primed dams were pooled from 6 mice per group. **(C-D)** Milk from PBS- and IAV-primed dams were pooled from 5 mice per group.

**Fig S10.**
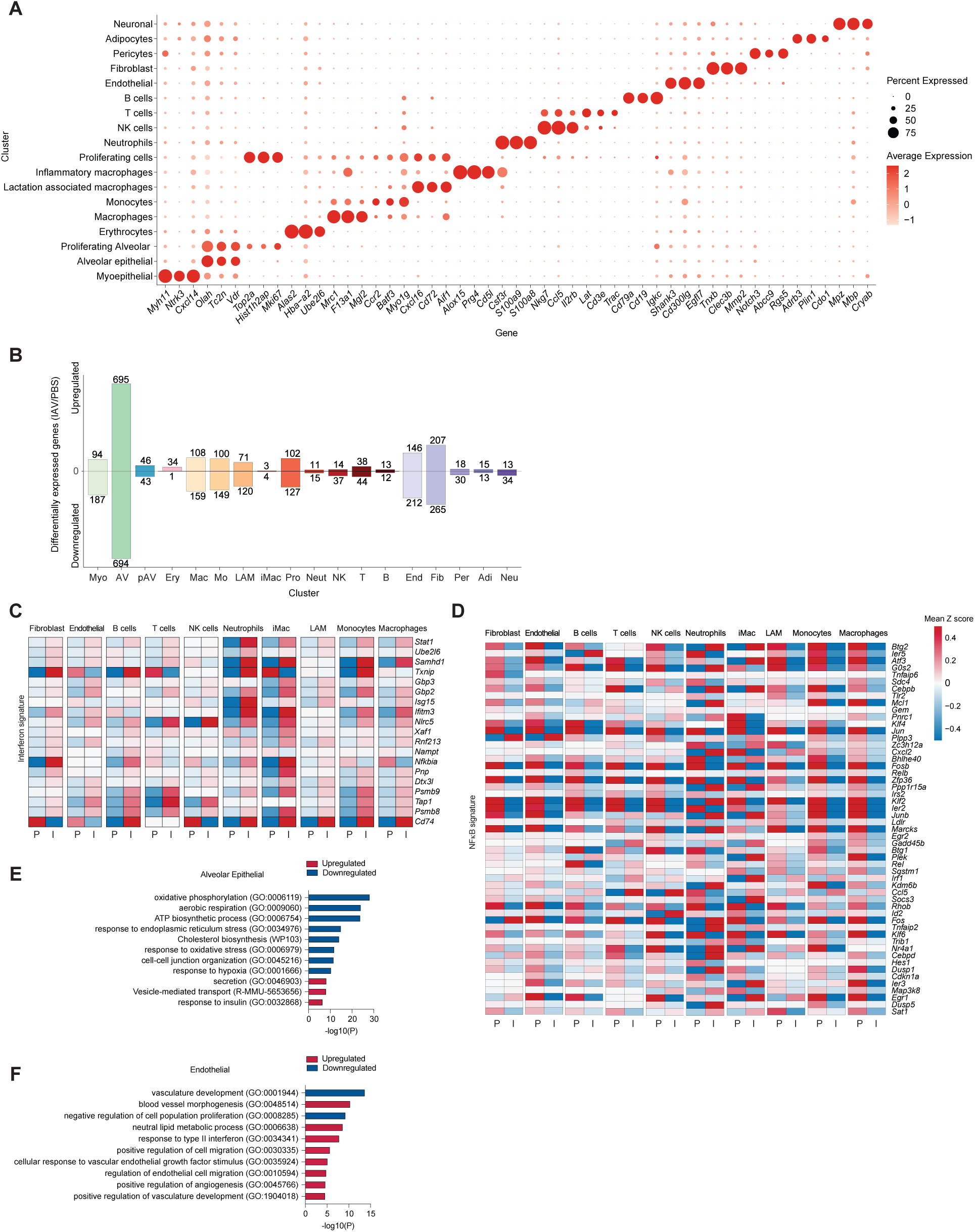
Preconceptual Respiratory Maternal IAV exposure induces long-term transcriptional changes in the mammary gland. Dams were infected with IAV (1 FFU, i.n.) or PBS and mated 21 days post infection. Cells from mammary glands of IAV and PBS-primed dams were analyzed by scRNA-seq at P5. **(A)** Dot plot depicting representative signature gene expressions across 18 distinct clusters. **(B)** Cells were analyzed for differential gene expression between conditions per cluster. Differential expression was determined by log2 fold change greater than 0.25 (upregulated) or less than −0.25 (downregulated), with padj<0.05 in IAV versus PBS condition. Number of differentially expressed genes across clusters in cells isolated from mammary glands of IAV-primed dams compared to those isolated from PBS-primed dams. Top bars represent upregulated genes, and bottom bars represent downregulated genes. **(C-D)** Heatmap of scaled expression of selected genes associated with interferon signature **(C)** and NF*κ*B signatures **(D)** in selected clusters. **(E-F)** Pathway enrichment analysis of differentially expressed genes showing significantly regulated biological processes for **(E)** alveolar epithelial cell clusters and **(F)** endothelial cell clusters. Differentially expressed genes with p adjusted value of <0.05 were used for analysis. See methods for scRNA-seq.

**Fig S11.**
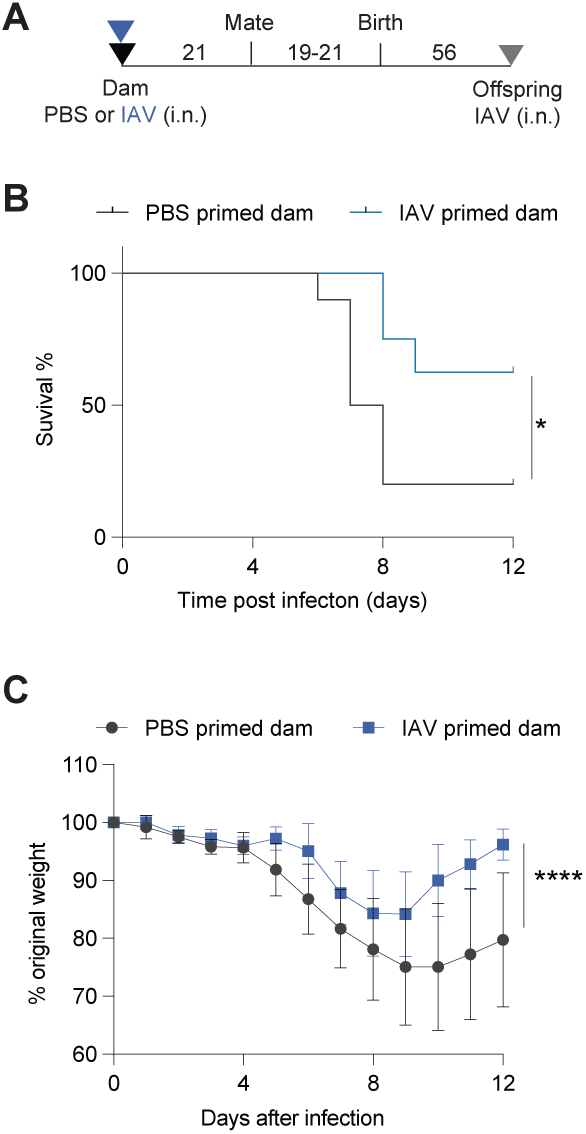
Maternal transferred immunity provides defense against lethal influenza infection in adult offspring. Dams were infected with IAV (1 FFU, i.n.) or PBS and mated 18-21 days post infection. Adult offspring of IAV or PBS primed dams were infected (10 FFU, IAV i.n.) at 8 weeks of age (P56). **(A)** Schematic of experimental setup. **(B)** Percent survival and **(C)** percent body weight change following intranasal IAV infection. Offspring of PBS-primed dam n =10; offspring of IAV-primed dam n=8. **(C)** Dot represents mean with error bars show SD. Statistical significance determined by survival Logrank test **(B)**, or 2-way ANOVA with fixed effects (type III) test, accounting time and experimental condition **(C)**. *p<0.05, **** p<0.0001.

